# ApoE4 impairs astrocyte lipid droplet formation via endolysosomal dysfunction

**DOI:** 10.64898/2026.09.01.748585

**Authors:** Isha Ralhan, Ju-Young Bae, Nathanael YJ Lee, Wendy Cai, Jinlan Chang, Suha Jarad, Hongmei Gu, Dawei Zhang, Maria S. Ioannou

## Abstract

An important component of neuron-glia coupling is the transport and delivery of neuron-derived lipids to glial lipid droplets. This pathway is important as failure to store incoming lipids in glial lipid droplets exacerbates neurodegeneration. ApoE4, a risk factor for Alzheimer’s disease, impairs this transport pathway. But how ApoE4 affects lipid trafficking in glia and whether these alterations can be restored is poorly understood. Here, we demonstrate that ApoE4 particles impair endolysosomal function and promote lipofuscin formation in cultured primary astrocytes, thereby disrupting the trafficking of lipids for storage into lipid droplets. Lipid droplets, however, can be recovered by preventing lysosomal impairment with PCSK9, a secreted protein that prevents LDLR recycling and reduces ApoE4 internalization. Lipid droplets are similarly recovered in the presence of ApoE4 by repairing endolysosomal function. Our findings reveal new insight into how ApoE4 dysregulates lipid storage while uncovering new mechanisms to correct these defects.

**Graphical Abstract:** 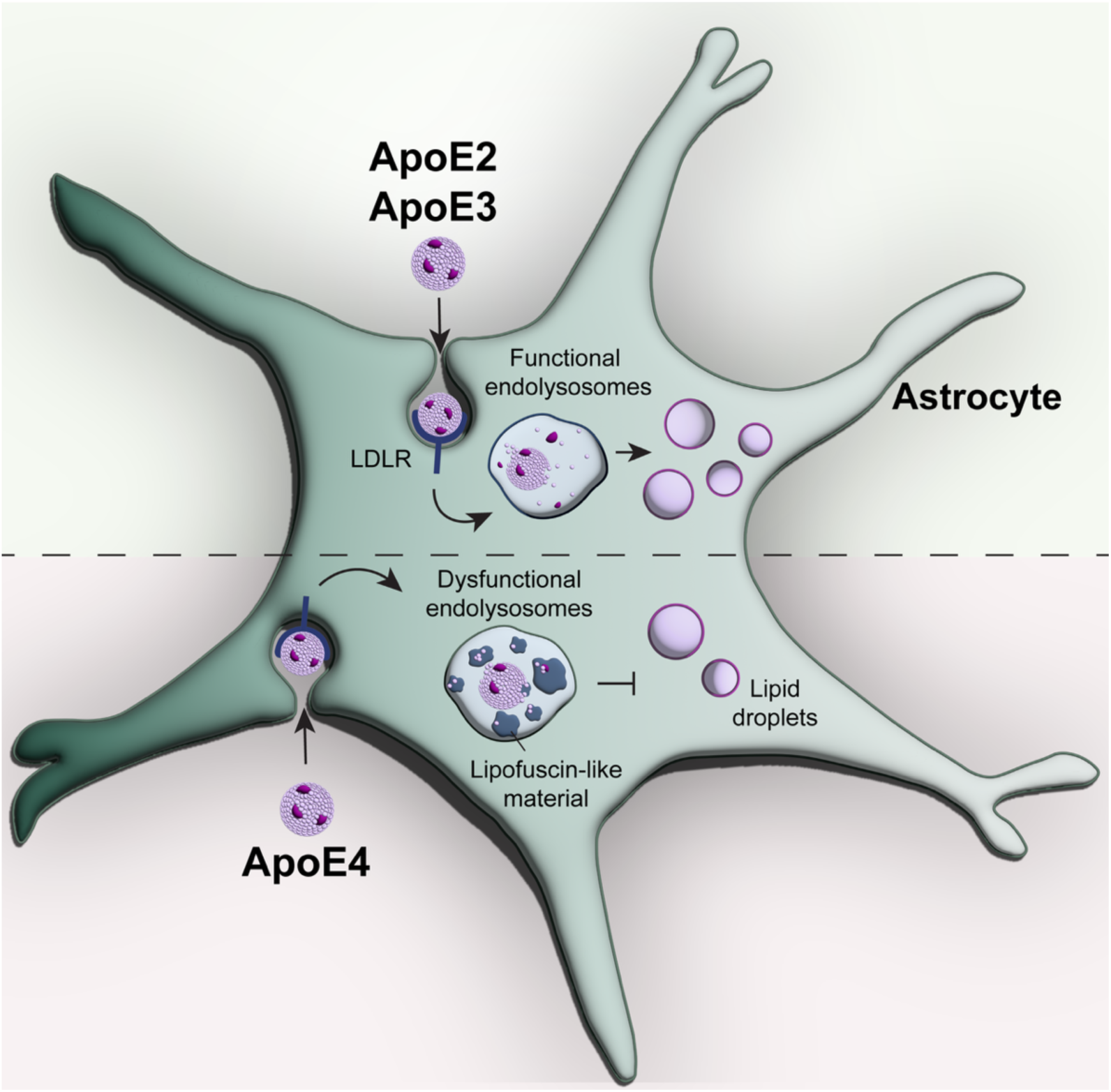

## INTRODUCTION

Astrocytes play an important role in regulating lipid homeostasis in the brain. During cellular stress, neurons rely on astrocytes to safely buffer and store excess lipids in lipid droplets. When lipid transport from neurons to astrocytes is impaired or if astrocytes fail to generate lipid droplets, neurons become vulnerable to degeneration^1–3^.

Apolipoprotein E (ApoE) is the most abundant apolipoprotein in the brain and is primarily expressed by astrocytes^4–7^. As a component of high-density lipoprotein-like particles, ApoE mediates the bidirectional lipid transport between neurons and glia. There are three common ApoE isoforms: ApoE2, ApoE3 and ApoE4, with the ApoE4 allele being a major risk factor for late-onset Alzheimer’s disease. Interestingly, knockdown of ApoE or expression of ApoE4 reduces the transfer of lipids to glial lipid droplets^1,2,8,9^. While this is, in part, due to reduced lipid efflux from neurons^10^, ApoE directly influences glial lipid trafficking and metabolism in an isoform-specific manner.

ApoE lipoproteins are internalized primarily by binding to members of the low-density lipoprotein receptor (LDLR) family^11^. Once internalized, ApoE lipoproteins are released from their cognate receptors and processed within the endolysosomal pathway. Within lysosomes, complex lipids originating from internalized lipoprotein particles as well as membranes derived from autophagy are degraded into simple lipids, such as fatty acids. The resulting simple lipids are transported out of lysosomes and if not used, converted to neutral lipids to be stored in lipid droplets^12^. Due to a reduced affinity for LDLR, the internalization of ApoE4 lipoproteins by glia is reduced relative to the other ApoE isoforms^13,14^. Despite reduced uptake, ApoE4 is still sufficient to induce endolysosomal defects^14,15^. But whether ApoE affects the storage of lipids in lipid droplets due to these endolysosomal defects and if restoring endolysosome function can recover lipid homeostasis is poorly understood.

Following endocytosis, LDLR dissociates from lipoprotein particles in the acidic pH of the endolysosome and is efficiently recycled back to the plasma membrane. In the periphery, LDLR can be engaged by the circulating proprotein convertase subtilisin/kexin type 9 (PCSK9), which reduces the half-life of LDLR by 90% by targeting it for degradation instead of recycling^16^. In doing so, PCSK9 dampens the uptake of ApoE lipoprotein particles. While PCSK9 is also expressed in the brain, its physiological role there is not well characterized^17^. In particular, whether PCSK9 affects lipid storage in glia remains unknown.

Here, we investigate how ApoE isoforms, as a component of extracellular lipid particles, affect astrocyte lipid homeostasis. We demonstrate that ApoE4 particles impair the transport of neuronal lipids to astrocytic lipid droplets, in part, via their isoform-specific effects on astrocytes. ApoE4 particles induce endolysosomal dysfunction, resulting in lipofuscin formation which disrupts fatty acid trafficking to lipid droplets. The endolysosomal defects and subsequent loss of lipid droplets can be rescued by extracellular PCSK9 which acts to limit ApoE4 internalization, or by reacidifying lysosomes. Altogether, this study reveals how ApoE4 impacts lipid droplet formation in astrocytes through its detrimental effects on the endolysosomal pathway.

## RESULTS

### ApoE genotype influences fatty acid transport from neurons to astrocytes

ApoE regulates lipid transport from neurons to glial lipid droplets^1,2^. To confirm the reported role of ApoE genotype in this process^2,8^, we cultured hippocampal neurons and glia from wild-type rats expressing rat ApoE3, or targeted replacement rats expressing human ApoE2, E3 and E4. We confirmed that over 95% of the glial cultures stained positive for astrocyte-specific glial fibrillary acidic protein (GFAP) indicating the enrichment of astrocytes in these cultures (Figures S1A-S1C). While 97% percent of neuronal cultures stained positive for the neuron-specific β3-tubulin (Figures S1D and S1E).

Neurons were loaded with BODIPY 558/568 (Red-C12), a fluorescently labelled saturated fatty acid. Red-C12 maintains fluorescence as it is incorporated into neutral lipids and phospholipids^18^, making it a useful reagent to track lipid transport. Neurons were then incubated together with unlabelled astrocytes on another coverslip separated by paraffin wax in Hanks’ Balanced Salt Solution (HBSS)^1,19^. The fluorescently labelled fatty acids are transferred from neurons to astrocytes in an ApoE-dependent fashion^1,8,10^. The appearance of neuron-derived Red-C12 in astrocytes was assessed by confocal microscopy (Figure 1A). We observed a genotype-specific effect whereby ApoE2 astrocytes had the highest amount of neuron-derived Red-C12 while ApoE4 astrocytes had the least, regardless of neuronal genotype (Figures 1B and 1C). These data confirm a genotype-specific role for ApoE in the transport of lipids from neurons to astrocyte lipid droplets.

**Figure 1.**
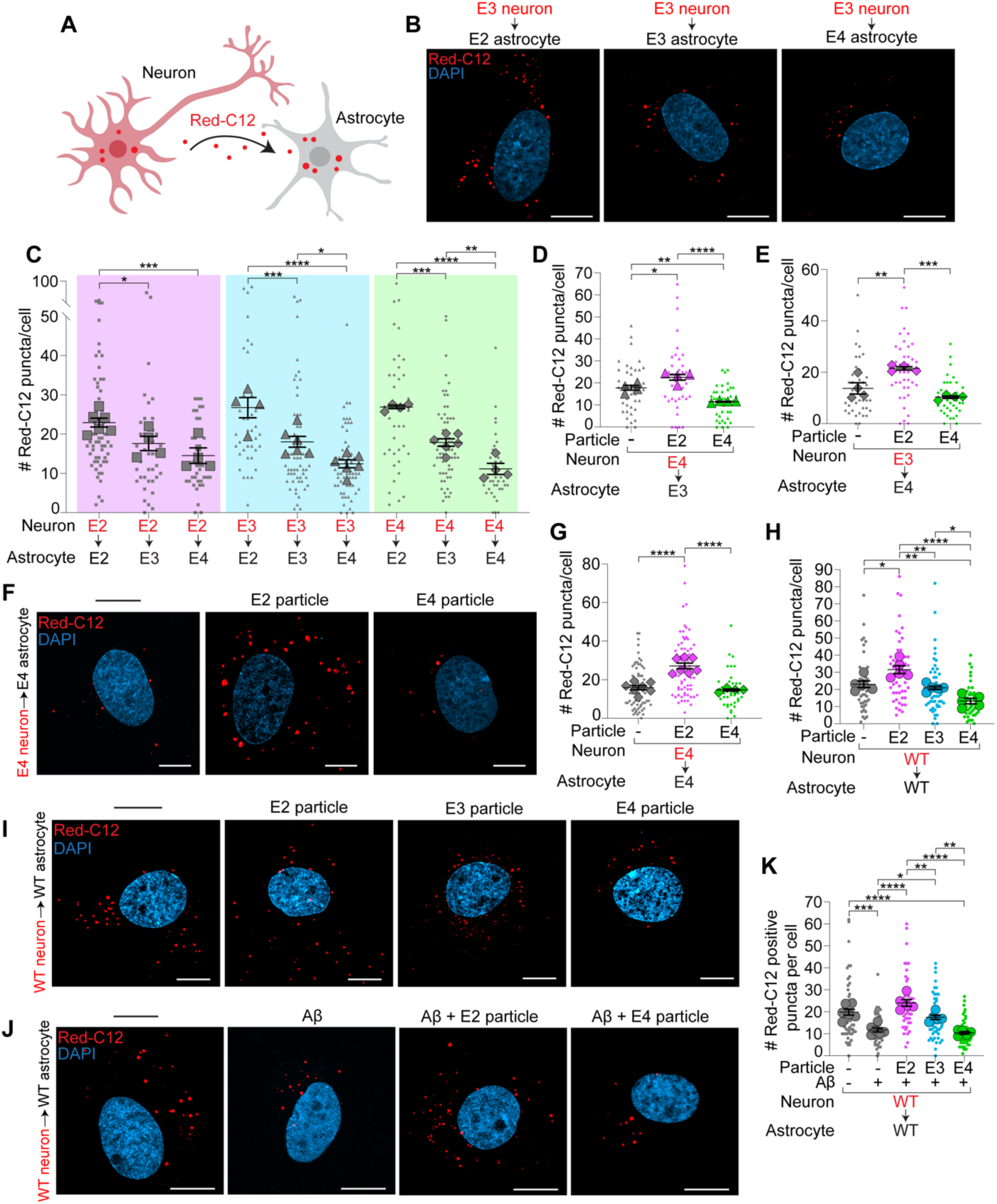
ApoE differentially regulates fatty acid transport from neurons to astrocytes. (A) Schematic of Red-C12 transfer assay from neurons to astrocytes. (B and C) Airyscan images of Red-C12 positive puncta in ApoE2, ApoE3 or ApoE4 astrocytes following fatty acid transfer assay in HBSS. n=4-6 independent experiments; mean ± SEM. Two-way ANOVA with Tukey’s post-test. Scale bars are 10 μm. (D) Red-C12 positive puncta in ApoE3 astrocytes following fatty acid transfer assay with ApoE4 neurons in HBSS ± 50 μg/ml ApoE2 or ApoE4 particles. n=4 independent experiments; mean ± SEM. One-way ANOVA with Tukey’s post-test. (E) Red-C12 positive puncta in ApoE4 astrocytes following fatty transfer assay with ApoE3 neurons in HBSS ± 50 μg/ml ApoE2 or ApoE4 particles. n = 4 independent experiments; mean ± SEM. One-way ANOVA with Tukey’s post-test. (F and G) Airyscan images of Red-C12 positive puncta in ApoE4 astrocytes following fatty acid transfer assay with ApoE4 neurons in HBSS ± 50 μg/ml ApoE2 or ApoE4 particles. n = 4-7 independent experiments; mean ± SEM. One-way ANOVA with Tukey’s post-test. Scale bars are 10 μm. (H) Red-C12 positive puncta in wild-type (WT) astrocytes following fatty acid transfer assay with WT neurons in HBSS ± 50 μg/ml ApoE2, ApoE3 or ApoE4 particles. n = 5 independent experiments; mean ± SEM. One-way ANOVA with Tukey’s post-test. (I) Airyscan images of WT astrocytes following transfer assay in HBSS ± 50 μg/ml ApoE2, ApoE3 or ApoE4 particles. Scale bars are 10 μm. (J and K) Airyscan images of WT astrocytes following Red-C12 transfer assay in media ± Aβ or 50 μg/ml ApoE2, ApoE3 or ApoE4 particles. n = 5-6 independent experiments; mean ± SEM. One-way ANOVA with Tukey’s post-test. Scale bars are 10 μm. All images displayed as maximum intensity projections. For all graphs independent replicates are in large shapes and technical replicates in small shapes; *p < 0.05, **p < 0.01, ***p < 0.001, ****p < 0.0001.

### Extracellular ApoE4 is sufficient to reduce fatty acid transport to astrocytes

Since ApoE genotype has a strong effect on lipid transfer to astrocytes, we next sought to investigate the role of ApoE isoforms on lipoprotein particles in the process. To explore this, we repeated the Red-C12 transfer assay in the presence of exogenous particles composed of 16:0-18:1 phosphatidylcholine and human ApoE2, ApoE3 or ApoE4^10^ (Figures S1F and S1G). The ApoE particles were approximately 10-20 nm in diameter as confirmed by transmission electron microscopy, with ApoE4 particles showing a slight but nonsignificant reduction in size compared to the other isoforms^20^ (Figures S1F and S1G). Addition of ApoE2 particles enhanced the amount of neuron-derived Red-C12 detected in ApoE3 and ApoE4 astrocytes (Figures 1D-1G). ApoE4 particles, on the other hand, failed to increase neuron-derived Red-C12 detected in astrocytes (Figures 1D-1G). In fact, ApoE4 particles decreased Red-C12 puncta detected in ApoE3 astrocytes (Figure 1D). Similar results were observed using wild-type neurons and astrocytes expressing rat ApoE3 (Figures 1H and 1I).

Since amyloid beta (Aβ) in the extracellular space increases in AD and correlates with disease progression^21,22^, we also tested how ApoE particles affect lipid transport in the presence of soluble Aβ42. Aβ42 reduced Red-C12 transport from wild-type neurons to astrocytes (Figures 1J and 1K). ApoE2 particles continued to enhance Red-C12 transport in the presence of Aβ42, while ApoE4 particles were unable to do so (Figures 1J, 1K, S1H, and S1I). These results point to an isoform-specific effect of ApoE isoforms on particles independent of the cell genotype in mediating neuron to astrocyte lipid trafficking.

### ApoE4 reduces lipid storage in astrocyte lipid droplets

We reasoned that the reduced neuron-derived Red-C12 detected in astrocytes in the presence of ApoE4 could be explained by several non-exclusive mechanisms. First, ApoE4 particles could efflux less lipids directly from neurons to deliver to astrocytes. Indeed, we previously found that ApoE4 particles extract less unsaturated phospholipids from neurons, however no difference in the amount of saturated Red-C12 efflux was observed^10^. This suggests that differences in the amount of Red-C12 trafficking from neurons is unlikely responsible for the effects observed here. We next assessed whether ApoE isoforms affect Red-C12 efflux from astrocytes, thereby differentially depleting astrocytes of the labelled lipids. Here, astrocytes were preloaded with Red-C12 and the Red-C12 released back into the media in the presence of particles was detected using a fluorescence plate reader. However, all ApoE particles promoted saturated Red-C12 efflux at similar levels (Figure 2A).

**Figure 2.**
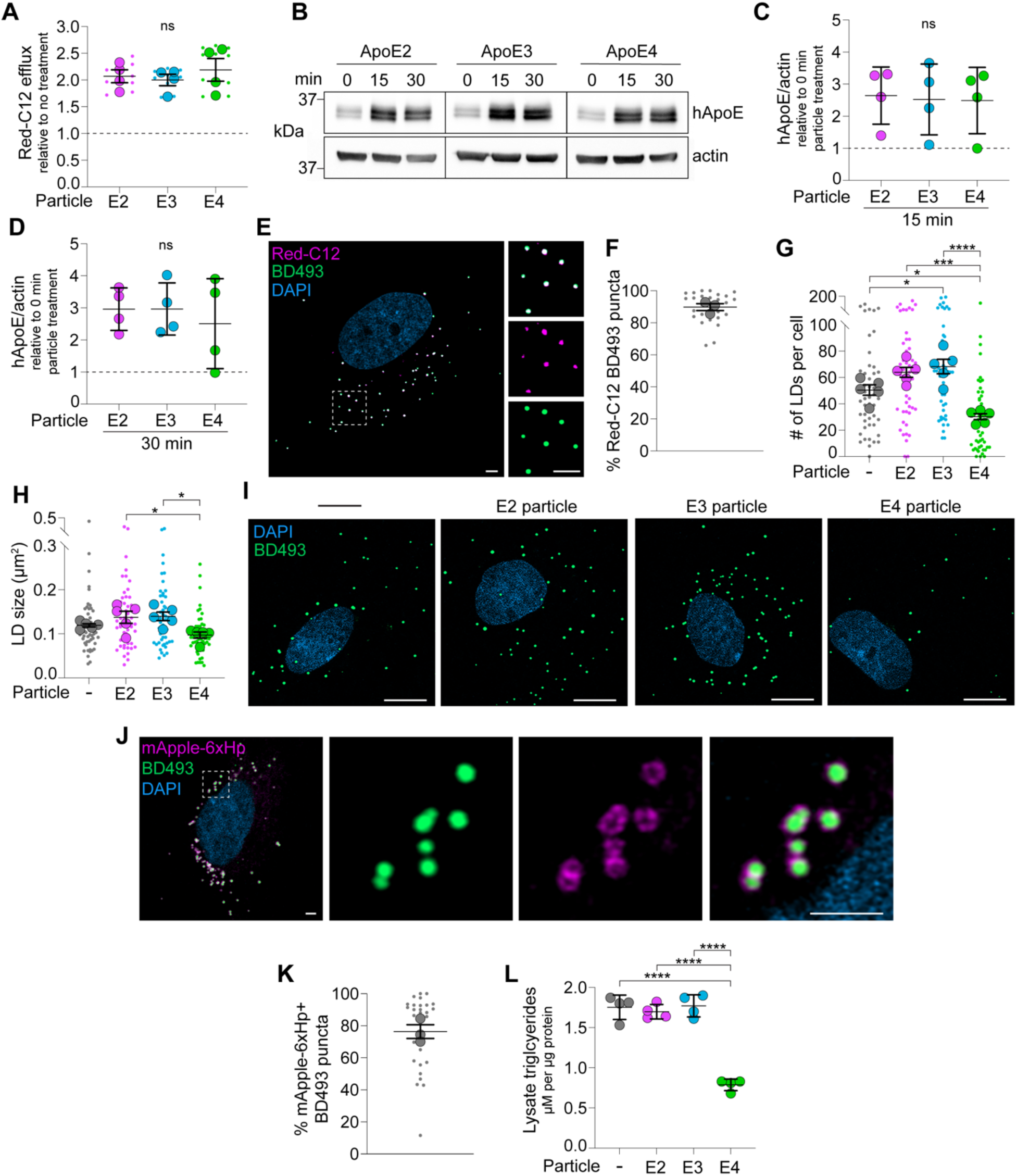
ApoE isoforms differentially regulate lipid storage in astrocytes. (A) Astrocytic-conditioned HBSS ± 50 μg/ml ApoE2, ApoE3 or ApoE4 particles analyzed for Red-C12 fluorescence and normalized to non-particle treated control neurons (dashed line). n = 4 independent replicates; mean ± SEM. One-way ANOVA with Tukey’s post-test. (B-D) Astrocytes ± 25 μg/ml ApoE2, ApoE3 or ApoE4 particles in HBSS were acid washed and analyzed by western blot for human ApoE (hApoE) and β-actin. 15- and 30-min particle treatments were normalized to 0 min particle treatment (dashed line). n = 4 independent replicates; mean ± SD. One-way ANOVA with Tukey’s post-test. (E and F) Airyscan image of astrocyte following Red-C12 transfer assay in HBSS. Boxed area magnified in right panels. The percentage of lipid droplets stained with BD493 containing Red-C12 were quantified. n = 3 independent replicates; mean ± SEM. Scale bars are 2.5 μM. (G and H) Quantification of BD493-positive lipid droplets (LDs) in astrocytes ± 50 μg/ml ApoE2, ApoE3 or ApoE4 particle treatment in HBSS. n = 5 independent experiments; mean ± SEM. One-way ANOVA with Tukey’s post-test. (I) Airyscan images of astrocytes displayed as maximum intensity projections ± 50 μg/ml ApoE2, ApoE3 or ApoE4 particle treatment in HBSS stained for BD493-positive lipid droplets. Scale bars are 10 μm. (J and K) Airyscan image of astrocyte showing mApple-6xHp and BD493. Boxed area magnified in right panels. The percent co-localization of mApple-6xHp and BD493 positive puncta were quantified. n = 3 independent replicates; mean ± SEM. Scale bars are 2.5 μM. (L) Astrocytes ± 50 μg/ml ApoE2, ApoE3 or ApoE4 particles in HBSS analyzed for total triglycerides and normalized to protein content. n = 4 independent experiments; mean ± SEM. One-way ANOVA with Tukey’s post-test. For all graphs independent replicates are in large shapes and technical replicates in small shapes; *p < 0.05, **p < 0.01, ***p < 0.001, ****p < 0.0001. ns, no significant differences were discovered.

Another possibility is that there are differences in the internalization of ApoE particles by astrocytes, as has been previously reported^14^. To test this, astrocytes were treated with particles in the absence of neurons, the extracellular pool was stripped by acid washing the cells, and the remaining intracellular pool of ApoE was analysed by western blot. We found that astrocytes internalized equivalent amounts of ApoE particles after 15 and 30 min of treatment, regardless of the isoform (Figures 2B-2D). While it remains possible that our assay is not sensitive enough to detect subtle differences in ApoE uptake, we reasoned that additional mechanisms likely play a role.

Since the Red-C12 fatty acids transferred to astrocytes accumulate in BODIPY-493/503 (BD493) positive puncta (Figures 2E and 2F), we wondered whether the differences in Red-C12 reflected changes in lipid droplet formation. In fact, storage of Red-C12 in lipid droplets is a requisite to detect Red-C12 puncta in these assays. To assess this, we evaluated BD493-positive lipid droplets in astrocytes after incubating with ApoE particles in the absence of neuronal lipids. We continued to perform these experiments in HBSS which is associated with increased lipid droplets through lysosomal degradation following autophagy^12^. Here, ApoE4 particles decreased the number and size of lipid droplets in wild-type astrocytes (Figures 2G-2I). Since ApoE4 particles did not influence the number or size of lipid droplets in ApoE4 astrocytes (Figures 2G-2I and S2A-S2C), we focused the remainder of our study on wild-type astrocytes to study the effects of ApoE isoforms on lipoprotein-like particles independent of cell genotype.

We also confirmed that these BD493 puncta are lipid droplets by co-localization with the lipid droplet marker mApple-6xHp^23^ (Figures 2J and 2K). Approximately 76% of the BD493 puncta co-localized with mApple-6xHp (Figures 2J and 2K). Since astrocytic lipid droplets are rich in triglycerides^24–26^, as a second strategy, we incubated astrocytes with particles and measured triglycerides directly. Consistent with our imaging data, we found that only ApoE4 particles decreased triglycerides in astrocytes (Figure 2L). Altogether, these results suggest that ApoE4 reduces lipid storage in lipid droplets under starvation conditions.

### ApoE4 reduces endolysosomal function and increases lipofuscin formation

Lysosomal degradation is an important source of lipids to be stored in lipid droplets^12,27^. Endolysosomal dysfunction is also prominent in glia expressing ApoE4 as well as in neurons treated with ApoE4 particles^10,14,15^. Therefore, one way that ApoE4 could affect lipid droplets is by disrupting the endolysosomal system. To assess late endosome-lysosomal compartments, we stained astrocytes with LysoTracker Red, a cell-permeable fluorescent dye that labels acidic compartments. We assessed whether ApoE particles differentially affect LysoTracker staining in astrocytes. We found that ApoE4 particles decreased the number of LysoTracker-positive puncta, while ApoE2 and ApoE3 particles had no effect (Figures 3A and 3B). LysoTracker puncta size was unaffected (Figure S3A). We then reasoned that if there were less acidic compartments, that endolysosomal function may be reduced. Indeed, astrocytes treated with ApoE4 particles had decreased activity of the endolysosomal protease cathepsin D (Figure 3C).

**Figure 3.**
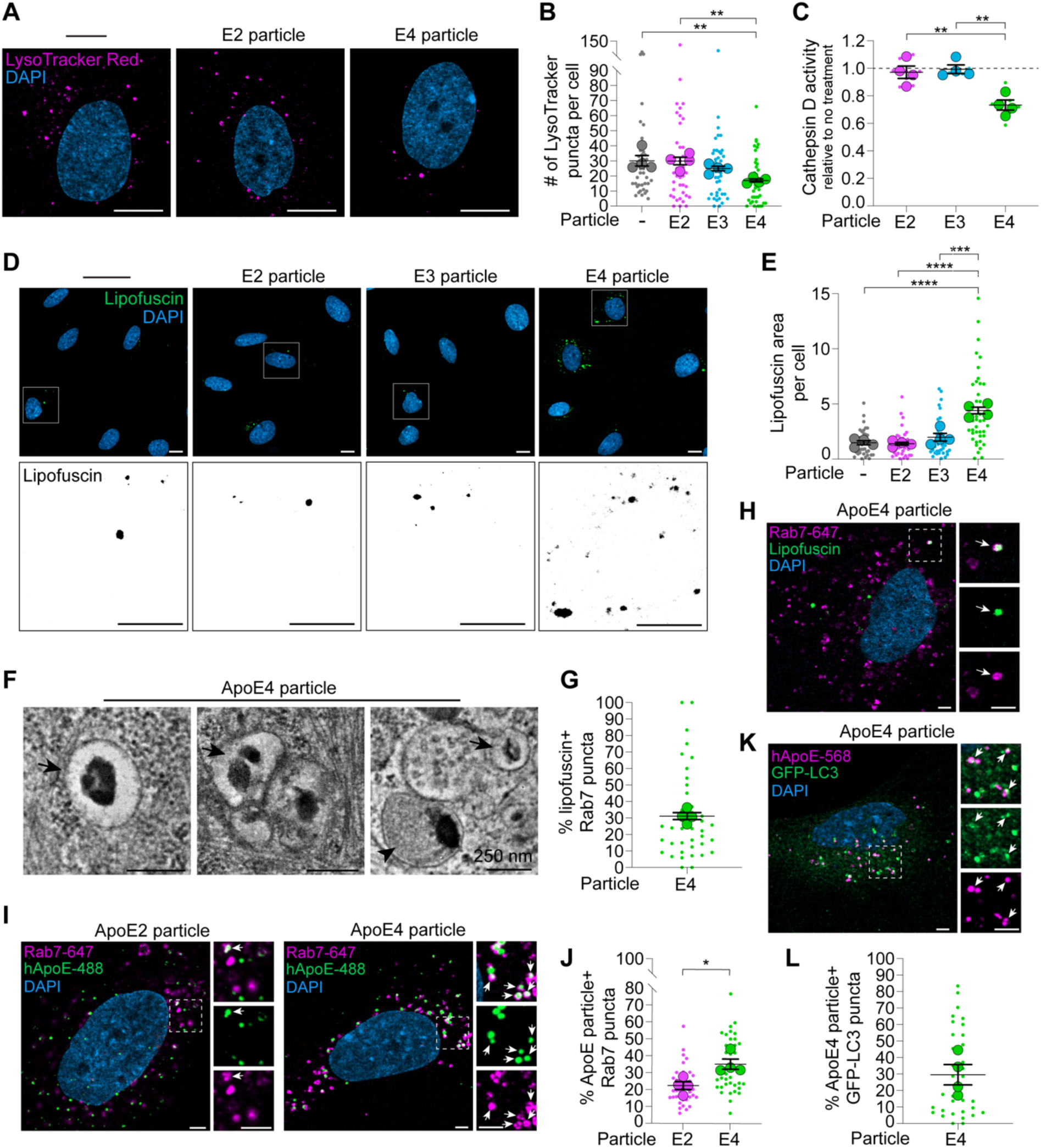
ApoE4 induces lipofuscin formation. (A) Airyscan images of astrocytes displayed as maximum intensity projections ± 50 μg/ml ApoE2 or ApoE4 particles in HBSS stained for LysoTracker Red. Scale bars are 10 μm. (B) Quantification of LysoTracker Red-positive puncta in astrocytes ± 50 μg/ml ApoE2, ApoE3 or ApoE4 particles in HBSS. n = 4 independent experiments; mean ± SEM. One-way ANOVA with Tukey’s post-test. (C) Astrocytes ± 50 μg/ml ApoE2, ApoE3 or ApoE4 particles in HBSS analyzed for cathepsin D activity and normalized to non-particle treated control neurons (dashed line). n = 6 independent experiments; mean ± SEM. One-way ANOVA with Tukey’s post-test. (D and E) Airyscan images displayed as maximum intensity projections of autofluorescent lipofuscin in astrocytes ± 50 μg/ml ApoE2, ApoE3 or ApoE4 particles in HBSS. Boxed area magnified in bottom panels. n = 4 independent experiments; mean ± SEM. One-way ANOVA with Tukey’s post-test. Scale bars are 10 μm. (F) Astrocytes treated with 50 μg/ml ApoE4 particles in HBSS stained with imidazole-buffered osmium were imaged by transmission electron microscopy. Arrows highlight lipofuscin-like material in electron-lucent endosomes. Arrowhead highlights lipofuscin-like material in electron-dense compartments. Scale bars are 250 nm. (G) Percent co-localization of lipofuscin-like autofluorescence and Rab7 puncta in astrocytes treated with ApoE4 particles in HBSS. n=4 independent experiments; mean ± SEM. (H) Airyscan image of astrocyte treated with 50 μg/ml ApoE4 particles in HBSS showing lipofuscin autofluorescence and Rab7. Boxed area magnified in right panels. Arrows highlight co-localization. Scale bars are 2.5 μM. (I and J) Airyscan image of astrocyte ± 50 μg/ml ApoE2 or ApoE4 particles in HBSS showing ApoE and Rab7. Boxed area magnified in right panels. Arrows highlight co-localization. The percent co-localization of human ApoE (hApoE) and Rab7 puncta were quantified. n = 4 independent replicates; mean ± SEM. Two-tailed Student’s t test. Scale bars are 2.5 μM. (K and L) Airyscan image of astrocyte treated with 50 μg/ml ApoE4 particles in HBSS showing GFP-LC3 and hApoE. Boxed area magnified in right panels. Arrows highlight co-localization. The percent co-localization of hApoE and GFP-LC3 puncta were quantified. n = 4 independent replicates; mean ± SEM. Scale bars are 2.5 μM. For all graphs independent replicates are in large shapes and technical replicates in small shapes; *p < 0.05, **p < 0.01, ***p < 0.001, ****p < 0.0001.

Lipofuscin is characterized by the accumulation of undigested material including oxidized lipids within non-degradative lysosomes^28^. Consistent with previous work^14^, we found that ApoE4, but not ApoE2 or ApoE3 particles, increased the number and area of autofluorescent puncta indicative of lipofuscin (Figures 3D, 3E and S3B). To further characterize this lipofuscin-like material we imaged astrocytes treated with ApoE4 particles using transmission electron microscopy. While electron-dense granules were found within many endosomal compartments, several of these endosomes appeared to be otherwise electron-lucent which is inconsistent with mature lipofuscin in lysosomes (Figure 3F). We reasoned that given the acute nature of our treatment, our imaging data likely includes late endosomes in early stages of lipofuscin formation^28^. Indeed, roughly 30% of the lipofuscin-like autofluorescent can be found co-localized with the late endosome marker Rab7 (Figures 3G and 3H).

Given that ApoE4 is disrupting endolysosomal function, we then assessed where internalized ApoE particles localize after the 4-hour treatment. We observed an increase in ApoE4 co-localization with Rab7-positive late endosomes as compared to ApoE2, accounting for roughly 35% of the ApoE4 puncta (Figures 3I and 3J). Approximately 20% of ApoE puncta co-localized with Rab5, a marker for early endosomes, though this was not affected by ApoE isoform (Figures S3C and S3D). Similarly, co-localization with LAMP1-positive lysosomes was unaffected by ApoE isoform, though this only accounted for less than 5% of the ApoE puncta, likely because its being degraded (Figures S3E and S3F). Interestingly, 30% of the ApoE4 puncta were also positive for GFP-LC3 (Figures 3K and 3L). This is consistent with the convergence of autophagic and endocytic-derived compartments^29^, and the ability of ApoE4 to disrupt autophagic degradation^30^. Collectively, this data suggests that internalized ApoE4 traffics through the endolysosomal pathway where it is predominately enriched in Rab7 late endosomes and that ApoE4 disrupts the function of endolysosomes and increases the formation of lipofuscin-like granules.

### PCSK9 prevents ApoE4-mediated lipofuscin formation and drop in lipid storage

Given the ability of ApoE4 to induce endolysosomal dysfunction, we reasoned that ApoE4 internalization would be required for its effects on lipid droplets. LDLR is a key receptor for ApoE-mediated uptake by astrocytes^13,14^. The activity of LDLR is tightly regulated by PCSK9, as it prevents its recycling to the surface and subsequently reduces ApoE uptake^31–33^. Here, we tested whether PCSK9 similarly reduces ApoE4 particle uptake in astrocytes. Astrocytes were pretreated with or without PCSK9 for 4 hours followed by ApoE4 particles for 15 minutes. The extracellular pool was stripped by acid washing the cells, and the remaining intracellular pool of ApoE was assessed by western blot. As predicted, PCSK9 effectively reduced ApoE4 particle internalization into astrocytes (Figures 4A and 4B). Since blocking ApoE4 endocytosis prevents lipofuscin formation^14^, we wondered if this would also be true with PCSK9 treatment. While PCSK9 alone had no effect on autofluorescent signals, it prevented the lipofuscin accumulation induced by ApoE4 particles (Figures 4C-4E). Finally, we tested if PCSK9 affects lipid droplets in ApoE4 treated cells. Again, PCSK9 alone had no effect on the number or size of BD493-positive lipid droplets, however, pretreatment with PCSK9 prevented the ApoE4-mediated reduction in lipid droplets in astrocytes (Figures 4F-4H). These results suggest that astrocytic lipid droplets are influenced by the uptake of ApoE and this can be regulated by PCSK9.

**Figure 4.**
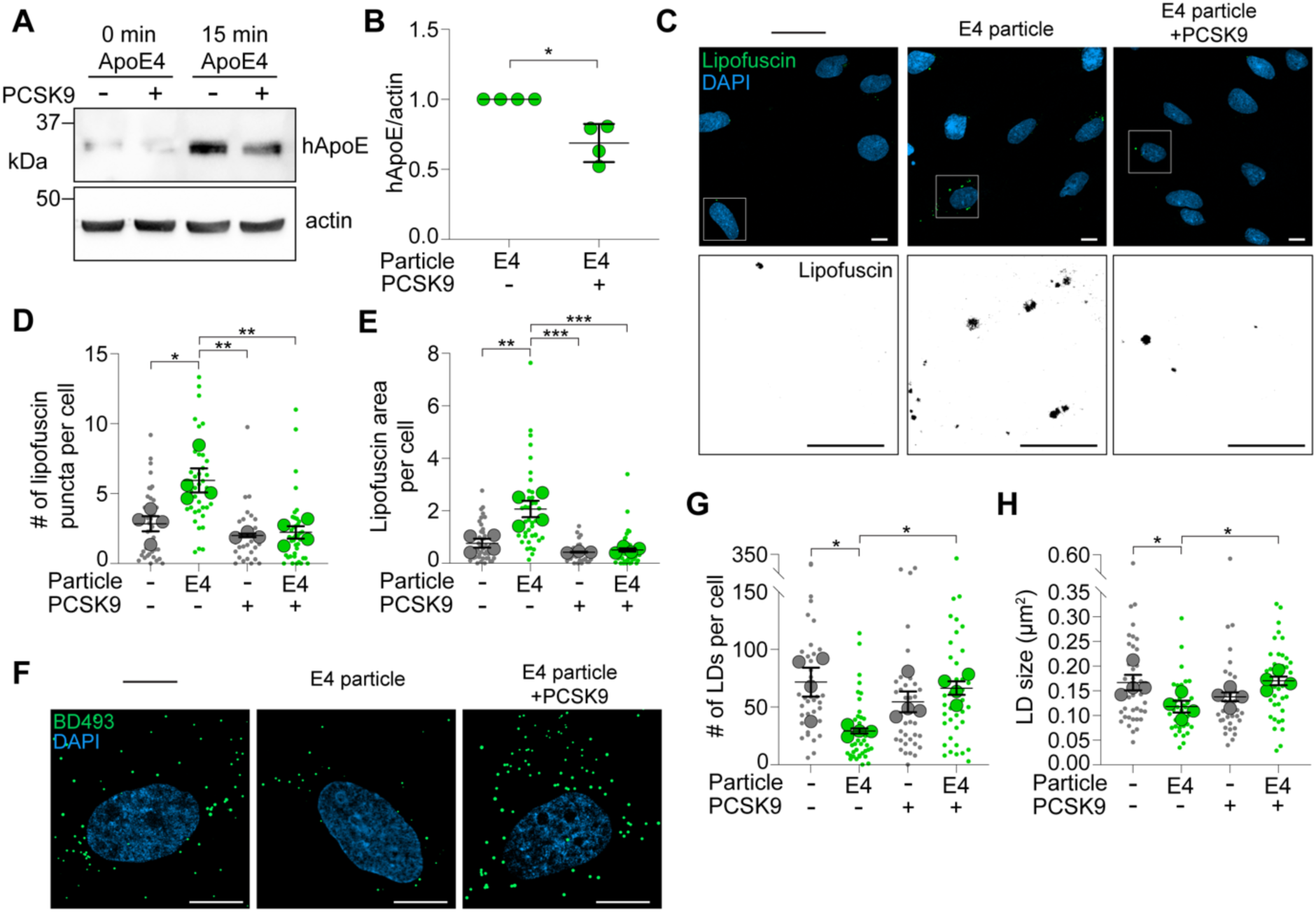
Reduced ApoE4 particle internalization restores astrocytic lipid droplet formation. (A) Astrocytes ± PCSK9 or 50 μg/ml ApoE4 particles in HBSS were acid washed and analyzed by western blot for human ApoE (hApoE) and β-actin. (B) Quantification of (A). Levels of hApoE/β-actin in PCSK9 treated astrocytes normalized to ApoE4 particle treated astrocytes following 15 min particle treatment. n = 4 independent replicates; mean ± SD. One sample t test with Bonferroni correction. (C-E) Airyscan images of autofluorescent lipofuscin in astrocytes ± PCSK9 or 50 μg/ml ApoE4 particles in HBSS. Boxed area magnified in bottom panels. n = 4 independent experiments; mean ± SEM. One-way ANOVA with Tukey’s post-test. Scale bars are 10 μm. (F-H) Airyscan images of astrocytes ± PCSK9 or 50 μg/ml ApoE4 particles in HBSS stained for BD493-positive lipid droplets (LDs). n = 4 independent replicates; mean ± SEM. One-way ANOVA with Tukey’s post-test. Scale bars are 10 μm. All images displayed as maximum intensity projections. For all graphs independent replicates are in large shapes and technical replicates in small shapes; *p < 0.05, **p < 0.01, ***p < 0.001, ****p < 0.0001.

### ApoE4 blocks lipid droplet formation through endolysosomal dysfunction

Given the requirement for ApoE internalization, we next explored whether dysregulated endolysosomes caused by ApoE4 are responsible for the effects on lipid droplets. First, we tested whether impairing endolysosomal function directly phenocopied ApoE4 treatment. Previous work found that starvation-induced lipid droplets are prevented by inhibiting lysosomal acidification and degradative function with bafilomycin A1^12,34^. We confirmed that bafilomycin A1 similarly reduced the number of LysoTracker-positive puncta (Figures S4A-S4C) and the number and size of lipid droplets in astrocytes (Figures S4D-S4F). Under these same conditions however, bafilomycin A1 had no effect on lipofuscin-like autofluorescence (Figures S4G-S4I). This supports the role of endolysosomal function in lipid droplet formation, independent of lipofuscin formation.

We next wondered if we could rescue these phenotypes by repairing lysosomal function. To assess this, we coarsely targeted two proton exchangers with known influence on lysosomal acidification. First, by inhibiting sodium-proton exchange (NHE). Pharmacological inhibition of NHE blocks proton efflux from the cell, thereby increasing cytosolic acidity. This could restore endolysosome acidity^35^, which has recently been reported^36,37^. One such inhibitor, rimeporide has previously been used to recover lysotracker staining and lysosome function^36,38–40^. Second, by stimulating the chloride-proton exchanger ClC-7^41–45^. Although ClC-7 imports two chloride ions in exchange for a proton, this displaces 3 positive charges which enhances the electrogenic proton pumping activity of the V-ATPase^43^. In some cases, the net result of increased chloride (and proton) import results in improved degradative luminal activity and clearance^41,43,44,46^. We indirectly stimulated ClC-7 by treating astrocytes with YM-201636 to selectively inhibit PIKfyve^47^. This serves to deplete PI(3,5)P_2_ levels which in turn relieves ClC-7 inhibition and hyper-activates the V-ATPase to lower endolysosomal pH^44,48^.

Here, we found that both rimeporide and YM-201636 rescued the ApoE4-mediated decrease in LysoTracker-positive puncta in astrocytes (Figures 5A-5D, S5A and S5B). Consistent with increased number of acidic vesicles, rimeporide and YM-201636 enhanced cathepsin D activity in the presence of ApoE4 particles (Figures 5E and 5F) and prevented the ApoE4-mediated formation of lipofuscin-like material (Figures 5G-5I, S5C and S5D). Finally, we reasoned that if lipid droplet formation in astrocytes is, in part, downstream of endolysosomal degradation, then improving lysosomal function should restore lipid droplet formation. Indeed, treatment with rimeporide or YM-201636 restored lipid droplet formation and triglyceride levels in astrocytes treated with ApoE4 particles (Figures 6A-6E, S5E and S5F). Collectively these results show that lipid droplet formation can be recovered in the presence of ApoE4 by restoring endolysosomal function.

**Figure 5.**
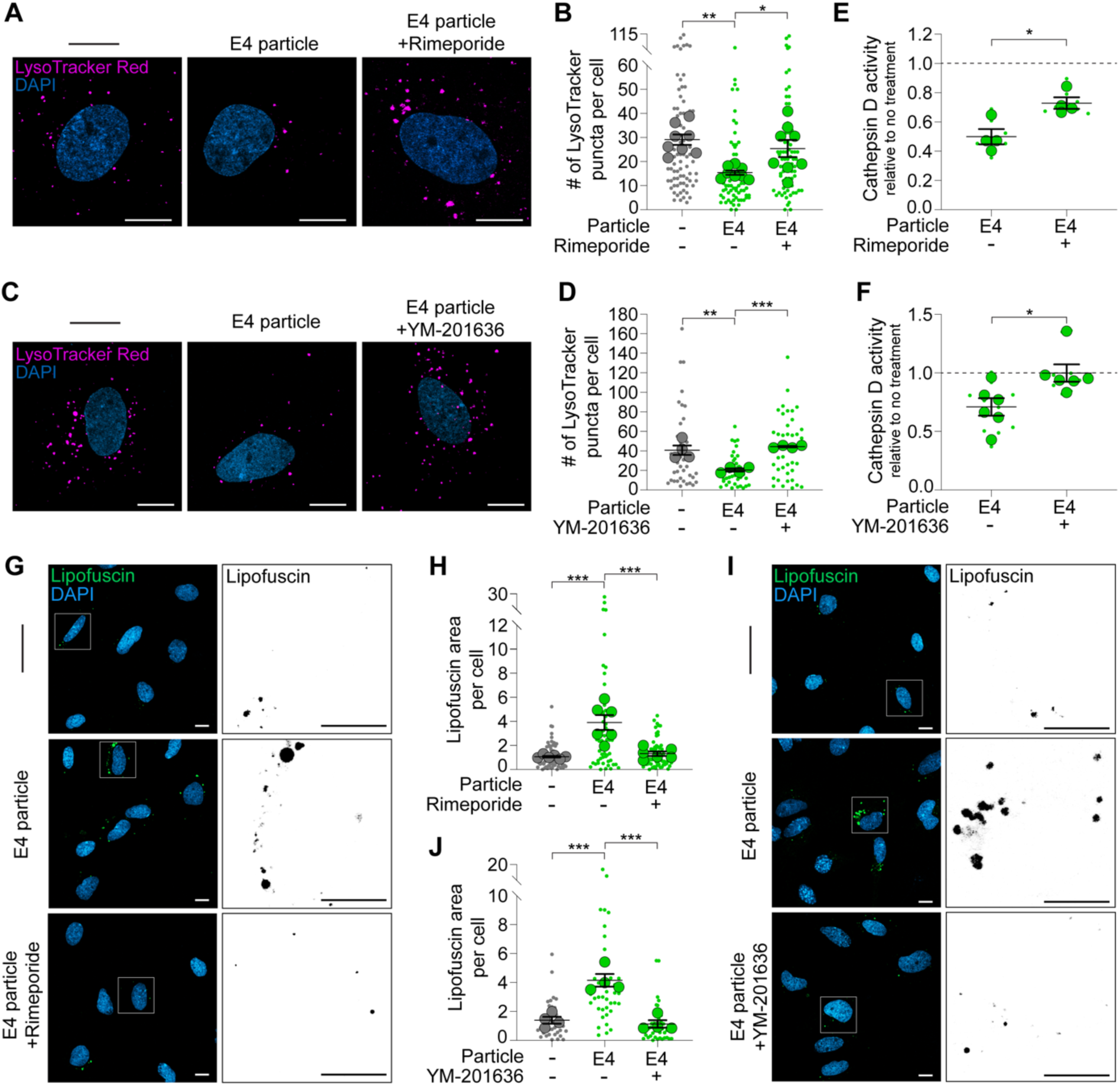
Restoring endolysosomal function prevents ApoE4-mediated lipofuscin accumulation. (A and B) Airyscan images of astrocytes ± rimeporide or 50 μg/ml ApoE4 particles in HBSS stained for LysoTracker Red. n = 8 independent experiments; mean ± SEM. One-way ANOVA with Tukey’s post-test. Scale bars are 10 μm. (C and D) Airyscan images of astrocytes ± YM-201636 or 50 μg/ml ApoE4 particles in HBSS stained for LysoTracker Red. n = 4 independent experiments; mean ± SEM. One-way ANOVA with Tukey’s post-test. Scale bars are 10 μm. (E) Astrocytes ± rimeporide or 50 μg/ml ApoE4 particles in HBSS analyzed for cathepsin D activity and normalized to non-particle treated control neurons (dashed line). n = 4 independent experiments; mean ± SEM. Two-tailed Student’s t test. (F) Astrocytes ± YM-201636 or 50 μg/ml ApoE4 particles in HBSS analyzed for cathepsin D activity and normalized to non-particle treated control neurons (dashed line). n = 6 independent experiments; mean ± SEM. Two-tailed Student’s t test. (G and H) Airyscan images of autofluorescent lipofuscin in astrocytes ± rimeporide or 50 μg/ml ApoE4 particles in HBSS. Boxed area magnified in right panels. n = 6 independent experiments; mean ± SEM. One-way ANOVA with Tukey’s post-test. Scale bars are 10 μm. (I and J) Airyscan images of autofluorescent lipofuscin in astrocytes ± YM-201636 or 50 μg/ml ApoE4 particles in HBSS. Boxed area magnified in right panels. n = 4 independent experiments; mean ± SEM. One-way ANOVA with Tukey’s post-test. Scale bars are 10 μm. All images displayed as maximum intensity projections. For all graphs independent replicates are in large shapes and technical replicates in small shapes; *p < 0.05, **p < 0.01, ***p < 0.001, ****p < 0.0001.

**Figure 6.**
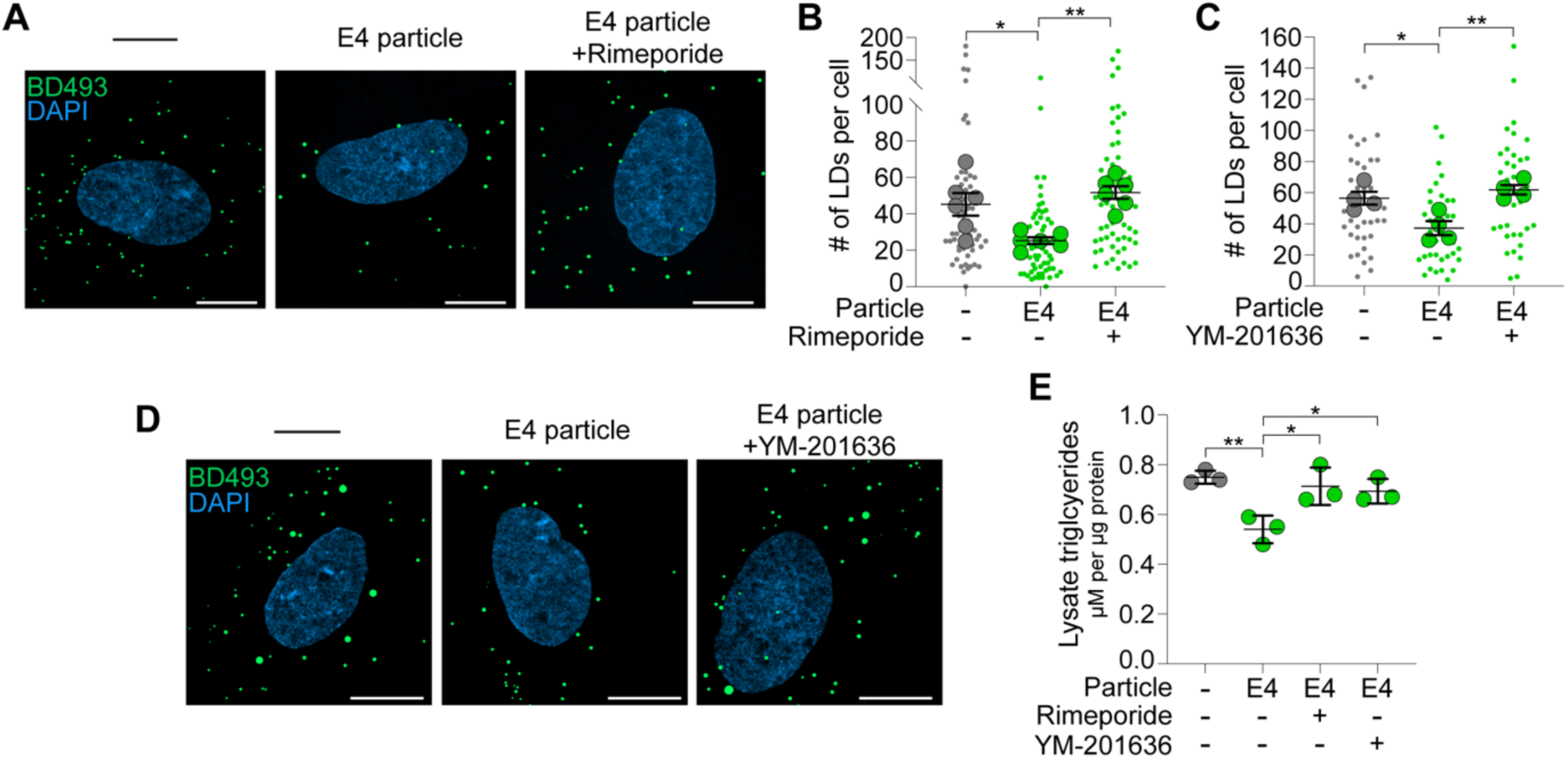
Restoring endolysosomal function recovers lipid droplet formation. (A and B) Airyscan images of astrocytes ± rimeporide or 50 μg/ml ApoE4 particle treatment in HBSS stained for BD493-positive lipid droplets (LDs). n = 6 independent experiments; mean ± SEM. One-way ANOVA with Tukey’s post-test. Scale bars are 10 μm. (C) Quantification of BD493-positive lipid droplets in astrocytes ± YM-201636 or 50 μg/ml ApoE4 particle treatment in HBSS. n = 4 independent experiments; mean ± SEM. One-way ANOVA with Tukey’s post-test. (D) Airyscan images of astrocytes ± YM-201636 or 50 μg/ml ApoE4 particle treatment in HBSS stained for BD493-positive lipid droplets. Scale bars are 10 μm. (E) Astrocytes ± rimeporide, YM-201636 or 50 μg/ml ApoE4 particles in HBSS analyzed for total triglycerides and normalized to protein content. n = 3 independent experiments; mean ± SEM. One-way ANOVA with Tukey’s post-test. All images displayed as maximum intensity projections. For all graphs independent replicates are in large shapes and technical replicates in small shapes; *p < 0.05, **p < 0.01, ***p < 0.001, ****p < 0.0001.

## DISCUSSION

Overall, this study uncovers new insight into how ApoE4 affects lipid homeostasis in the brain. We show that uptake of ApoE4 on lipoprotein-like particles impairs endolysosomal function. This prevents the degradation of lipids resulting in lipofuscin formation and limits the supply of lipids for storage in lipid droplets. Finally, preventing ApoE uptake as well as restoring lysosomal activity can effectively recover lipid droplet formation.

ApoE has well documented roles on lipid droplet physiology. Astrocytes expressing ApoE4 typically have more lipid droplets compared to those expressing ApoE2 or ApoE3^49–53^. This is due to altered lipid metabolism in ApoE4 astrocytes, such as reduced fatty acid oxidation, which increases the need for lipid storage^8^. Lipid droplets in ApoE4 astrocytes are also resistant to degradation due to impaired autophagy^50,54^. Importantly, these studies were performed on astrocytes under nutrient-rich conditions. In addition, treating astrocytes with particles allowed us to probe the effects of internalized ApoE independent of ApoE genotype. This is particularly important given the newly discovered role of ApoE in escaping the secretory pathway to directly influence lipid droplets^50^. We also performed our experiments under starvation which is a robust method to increase lipid droplet formation using lipids sourced from autophagy^12^. Under these conditions, as opposed to increased lipid droplets, we find the opposite to be true, that ApoE4 reduced the formation of lipid droplets. This is likely because under starvation a major determinant of lipid storage is lysosomal degradation following autophagy which is disrupted by ApoE4. We speculate that under physiological conditions, both processes, lipid droplet growth and degradation, are affected by ApoE isoform.

In addition to autophagy supplying lipids to growing lipid droplets, astrocytes also take up and store lipids released by neurons. This pathway protects neurons from toxicity associated with oxidized lipids, as failure to store neuronal lipids in glial lipid droplets results in neurodegeneration. In this scenario, lipid droplets are thought to protect cells from lipid peroxidation and ferroptosis^26^. Though it is still unclear whether this is by sequestering oxidized lipids to limit propagation of lipid peroxidation or alternatively, whether they store unmodified polyunsaturated fatty acids to protect them from becoming peroxidated in the first place. Regardless, failure to regulate lipid peroxidation can trigger ferroptosis, a form of cell death driven by the iron-dependent accumulation of lipid peroxides^55^.

Since ApoE plays a key role in several steps of this neuron-to-glia transport pathway, it is not surprising that multiple steps are influenced by ApoE isoforms. For example, in *Drosophila*, glial lipid droplets that form in response to oxidative stress in neurons, are blocked by knocking out the apolipoprotein Lazarillo^2^. This can be rescued by expressing ApoE2 but not ApoE4^2^. Consistent with this, ApoE4 is less efficient at effluxing oxidized lipids from neurons as compared to the protective isoforms ApoE2 or ApoE3 containing the Christchurch mutation^10^. ApoE4 is also less efficient at being internalized by astrocytes through LDLR^14^. While this appears contrary to previous studies showing increased LDLR binding by ApoE4 relative to the other isoforms^56–58^, these results may be influenced by the lipidation status of ApoE, the types of lipids bound and even the post-translational modifications of ApoE^59^. Here, we were unable to detect isoform-dependent differences in ApoE uptake by astrocytes, however, we did confirm that internalization of ApoE4 is critical for its effects on lipid droplets. We found that treating astrocytes with the LDLR modulator PCSK9 can restore glial lipid droplet formation. PCSK9 is a secreted factor that binds to LDLR and promotes its trafficking away from recycling towards degradation. It is also known to target other members of the LDLR family for degradation, such as, very low-density lipoprotein receptor (VLDLR) and Apolipoprotein E receptor 2 (ApoER2)^60–63^. As such, the effects observed in this study may not be specific to just LDLR. For example, LRP1 is also a risk factor for Alzheimer’s disease and disruption of LRP1 reduces glial lipid droplet formation^64^. Interestingly PCSK9 can also target LRP1 for degradation^63,65^. This supports the involvement of ApoE receptors in addition to LDLR in this process.

Our findings also reveal that internalization of ApoE4 impairs lysosomal function leading to an accumulation of lipofuscin-like material. This supports recent work demonstrating that ApoE4 carrying polyunsaturated cholesterol esters induced lysosomal defects and lipofuscin accumulation^14^. Lipofuscin is a pathological autofluorescent lipopigment characterized by the accumulation of partially digested oxidized proteins, lipids and metals within lysosomes^28^. While lipofuscin accumulation is commonly observed in neurons during aging and in neurodegenerative disease^66–69^, it has also been found in microglial lysosomes of young brains^70^, suggesting its accumulation can be progressive and reflect early lysosomal pathology. Here, we assessed lipofuscin-like autofluorescence as a proxy for oxidized lipid accumulation that could be resolved when lysosomal function was improved. An important question that remains is at what stage does lipofuscin become indigestible? Given that a large proportion of the lipofuscin-like granules are within Rab7-positive late endosomes, this could reflect early stages of lipofuscin that can still be degraded. However, if left unresolved, it develops into an indigestible form associated with disease^28^. This lipofuscin contributes to production of reactive oxygen species^71^ and can damage lysosomal membranes^71^, further enhancing cellular stress. The reciprocal relationship between lysosomal dysfunction and lipofuscin deposition generates a vicious cycle that can further exacerbate neurodegeneration.

Here we report potential ways to break the cycle; by restoring endolysosomal function, lipofuscin formation can be avoided as lipids are degraded and stored safely in lipid droplets. This was accomplished by targeting solute transporters in the cell, particularly NHE-1 and albeit indirectly ClC-7. While stimulating ClC-7 activity with PIKfyve inhibition improves lysosomal function^41,43,44,48^, caution should be exercised with this approach as unfavourable effects on autophagy and lysosomal function have also been reported^72–74^. Ideally, targeting ClC-7 directly could be used to improve lysosomal function without the broad effects of inhibiting PIKfyve. As an alternative strategy we also inhibited NHE-1 to increase endolysosomal acidity and function. These results are consistent with studies showing NHE-1 inhibition restores lysosome function in models of Parkinson’s disease^36,40^. In fact, astrocyte-specific knockdown of NHE-1 improved outcomes such as hypertrophy and gliosis in models of ischemic stroke^75^. This is likely due to improved lysosomal function, lipid degradation and storage. While effective, targeting NHE-1 is also not specific to endolysosomes as NHE-1 is most abundant at the plasma membrane^76,77^. However, inhibition of lysosome-specific NHEs have shown similar promise for improving cellular function in models of neurodegenerative disease. For example, NHE-6 depletion reduced ApoE4-mediated amyloid-plaque load^39^. Altogether our work continues to support the potential of restoring lysosomal function as a protective mechanism for neurodegenerative disease.

Finally, a proposed function of glial lipid droplet formation is to protect against oxidized lipids derived from neurons^2,64^. Although glial lipid droplets are initially neuroprotective^1,2^, their capacity to manage sustained lipid burden remains unclear. For example, excessive lipid droplet formation in glia is linked to inflammation during aging and disease. This was shown in the aging brain and in models of tauopathy, where lipid droplet laden microglia exhibit a dysfunctional, proinflammatory phenotype characterized by reduced phagocytic capacity, increased reactive oxygen species production and enhanced secretion of proinflammatory cytokines^78–80^. Yet failure to form lipid droplets in microglia, for example by acyl-CoA diacylglycerol acyltransferase (DGAT) knockout, exacerbates neurodegeneration^81^. Lipid droplets in astrocytes are similarly associated with changes in lipid metabolism and inflammation^82–84^. We speculate that under physiological conditions, glia have the capacity to buffer neuron-derived lipids. However, when the influx of lipids exceeds their capacity to neutralize, degrade or store these lipids, glial function can become compromised, contributing to neurodegeneration.

## RESOURCE AVAILABILITY

### Lead contact

Further information and requests for resources and reagents should be directed to and will be fulfilled by the lead contact, Maria S. Ioannou.

### Materials availability

This study did not generate new unique reagents.

### Data and code availability

- All data reported in this paper will be shared by the lead contact upon request.
- This paper does not report original code.
- Any additional information required to reanalyze the data reported in this paper is available from the lead contact upon request.

## ACKNOWLEDGMENTS

We thank Spencer Freeman for valuable advice on lysosomes and helpful comments on the manuscript. We thank Pinzhang Gao and Suey Van Baarle for assistance with electron microscopy sample preparation. Experiments were performed at the University of Alberta Faculty of Medicine & Dentistry Cell Imaging Core, RRID:SCR_019200. I.R. was supported by a Canada Graduate Scholarship from the Canadian Institutes of Health Research Doctoral Award #181551 and an Izaak Walton Killam Memorial Scholarship. M.S.I. is supported by grants from the Canadian Institutes of Health Research (#173321), Natural Sciences and Engineering Research Council of Canada (#2020-04047), and the Canadian Research Chairs Program (#2021-00027).

## AUTHOR CONTRIBUTIONS

Conceptualization, I.R. and M.S.I.; Investigation, I.R., J.B., and W.C.; Analysis, I.R., J.B., N.Y.J.L., and W.C.; Resources, J.C., S.J., H.G., and D.Z.; Writing – Original Draft, I.R. and M.S.I.; Writing – Review & Editing, I.R., J.B., N.Y.J.L., W.C., J.C., S.J., H.G., D.Z., and M.S.I.; Supervision, D.Z. and M.S.I.

## STAR METHODS

### KEY RESOURCES TABLE

| REAGENT or RESOURCE | SOURCE | IDENTIFIER |
| --- | --- | --- |
| Antibodies |  |  |
| Mouse monoclonal anti-β3-tubulin | Biolegend | Cat# 801202;<br>RRID:AB_2313773 |
| Rabbit polyclonal anti-Iba1 | Wako | Cat# 019-19741;<br>RRID:AB_839504 |
| Chicken polyclonal anti-GFAP | Abcam | Cat# ab4674;<br>RRID:AB_304558 |
| Goat polyclonal anti-Apolipoprotein E | Millipore-Sigma | Cat# AB947;<br>RRID:AB_2258475 |
| Mouse monoclonal anti-β-actin | Cell Signaling Technology | Cat# 3700;<br>RRID:AB_2242334 |
| Rabbit polyclonal anti-LAMP1 | Abcam | Cat# ab24170;<br>RRID:AB_775978 |
| Mouse monoclonal anti-Rab5A [E6N8S] | Cell Signalling Technology | Cat# 46449;<br>RRID:AB_2799303 |
| Rabbit monoclonal anti-Rab7 [EPR7589] | Abcam | Cat# ab137029;<br>RRID:AB_2629474 |
| Goat anti-Mouse IgG (H+L) Cross-Adsorbed, Alexa Fluor 488 | Thermo Fisher Scientific | Cat# A-11001;<br>RRID:AB_2534069 |
| Goat anti-Rabbit IgG (H+L) Cross-Adsorbed, Alexa Fluor 488 | Thermo Fisher Scientific | Cat# A-11008;<br>RRID:AB_143165 |
| Donkey anti-Goat IgG (H+L) Cross-Adsorbed, Alexa Fluor 488 | Thermo Fisher Scientific | Cat# A-11055;<br>RRID:AB_2534102 |
| Donkey anti-Goat IgG (H+L) Cross-Adsorbed, Alexa Fluor 568 | Thermo Fisher Scientific | Cat# A-11057;<br>RRID:AB_2534104 |
| Donkey anti-Rabbit IgG (H+L) Cross-Adsorbed, Alexa Fluor 647 | Thermo Fisher Scientific | Cat# A-31573;<br>RRID:AB_2536183 |
| Goat anti-Mouse IgG (H+L) Cross-Adsorbed, Alexa Fluor 647 | Thermo Fisher Scientific | Cat# A-21235;<br>RRID:AB_2535804 |
| Goat anti-Chicken IgY (H+L), Alexa Fluor 647 | Thermo Fisher Scientific | Cat# A-21449;<br>RRID:AB_2535866 |
| Rabbit anti-Goat IgG (H+L), HRP | Thermo Fisher Scientific | Cat# 31402;<br>RRID:AB_228395 |
| Peroxidase-AffiniPure Donkey anti-Mouse IgG | Jackson ImmunoResearch Labs | Cat# 715-035-150;<br>RRID:AB_2340770 |
| Chemicals, peptides, and recombinant proteins |  |  |
| ApoE2 POPC | Kisbee Therapeutics | N/A |
| ApoE3 POPC | Kisbee Therapeutics | N/A |
| ApoE4 POPC | Kisbee Therapeutics | N/A |
| POPC | Avanti Polar Lipids | Cat# 850457C |
| Sodium cholate | Sigma-Aldrich | Cat# C6445 |
| Cytosine-beta-D-arabinofuranose hydrochloride (AraC) | Thermo Fisher Scientific | Cat# J6567103 |
| BODIPY 558/568 C12 (4,4-Difluoro-5-(2-Thienyl)- 4-Bora-3a,4a-Diaza-s-Indacene-3-Dodecanoic Acid) (Red-C12) | Thermo Fisher Scientific | Cat# D3835 |
| CholEsteryl BODIPY FL C12 (4,4-Difluoro-5,7-Dimethyl-4-Bora-3a,4a-Diaza-s-Indacene-3-Dodecanoate (BD-CE) | Thermo Fisher Scientific | Cat# C3927MP |
| BODIPY 493/503 (BD493) | Thermo Fisher Scientific | Cat# D3922 |
| DAPI Staining Solution | Abcam | Cat# ab228549 |
| β-Amyloid 1-42, aβ, ultra pure, TFA, recombinant human | Millipore-Sigma | Cat# AG912 |
| LysoTracker Red DND-99 | Thermo Fisher Scientific | Cat# L7528 |
| Bafilomycin A1 | Sigma-Aldrich | Cat# B1793 |
| Rimeporide (EMD-87580) | Ray Biotech | Cat# 332-12602-1 |
| YM-201636 | Cayman Chemical | Cat# 13576 |
| Halt Protease and Phosphatase Inhibitor Cocktail | Thermo Fisher Scientific | Cat# 78440 |
| Lipofectamine 3000 Transfection Reagent | Thermo Fisher Scientific | Cat# L3000015 |
| Dimethyl sulfoxide | Sigma-Aldrich | Cat# 472301; CAS: 67-68-5 |
| Critical commercial assays |  |  |
| Cathepsin D Activity Assay Kit | Abcam | Cat# ab65302 |
| Pierce BCA Protein Assay Kit | Thermo Fisher Scientific | Cat# 23227 |
| Triglyceride-Glo Assay Kit | Promega | Cat# J3160 |
| Experimental models: Organisms/strains |  |  |
| Rat: Sprague Dawley Wild type | Charles River | CD(SD): 001 |
| Rat: Sprague Dawley HAPOE2 | Envigo | 39402 |
| Rat: Sprague Dawley HAPOE3 | Envigo | 39502 |
| Rat: Sprague Dawley HAPOE4 | Envigo | 35902 |
| Recombinant DNA |  |  |
| mApple-6xHp | Li et al, 2024 | Addgene plasmid #214410 |
| pEGFP-LC3 | Hailey et al, 2010 | N/A |
| Software and algorithms |  |  |
| (Fiji is just) ImageJ 2.1.0 | NIH | https://imagej.net/; RRID:SCR_003070 |
| Graphpad Prism 10 | Graphpad | https://www.graphpad.com/; RRID:SCR_002798 |
| Microsoft Excel 365 | Microsoft Corporation | https://office.com/ |
| ZEN Software | Zeiss | https://zeiss.ca/ |
| Bio Tek Synergy H1 Multimode Reader | Agilent | https://www.agilent.com |
| Adobe Illustrator | Adobe | https://www.adobe.com; RRID:SCR_010279 |
| Other |  |  |
| Basal Medium Eagle (BME) | GIBCO | Cat# 21010046 |
| Neurobasal Medium | GIBCO | Cat# 21103049 |
| Dulbecco's Modified Eagle's Medium (DMEM) high glucose | Sigma-Aldrich | Cat# D5796 |
| B-27 Supplement (50X), serum free | GIBCO | Cat# 17504044 |
| Antibiotic-Antimycotic | GIBCO | Cat# 15240062 |
| Fetal bovine serum | GIBCO | Cat# 12483020 |
| Sodium pyruvate | GIBCO | Cat# 11360070 |
| GlutaMAX | GIBCO | Cat# 35050061 |
| HEPES | GIBCO | Cat# 15630080 |
| HBSS without Ca<sup>2+</sup> and Mg<sup>2+</sup> (for dissections) | Cytiva | Cat# SH30588.01 |
| HBSS with Ca<sup>2+</sup> and Mg<sup>2+</sup> (for experiments) | Cytiva | Cat# SH30268.01 |
| DPBS/Modified | Cytiva | Cat# SH30028.03 |
| Poly-D-lysine Hydrobromide | Sigma-Aldrich | Cat# P6407 |
| Papain Dissociation Enzyme | Worthington Biochemical | Cat# LK003176 |
| Benzonase Nuclease | Millipore-Sigma | Cat# E1014 |
| DAKO Fluorescence Mounting Media | Agilent Technologies | Cat# S3023 |
| 25-mm round cover glass coverslips (#1.5) | Harvard Apparatus Canada | Cat# 640715 |
| ANTI-FLAG M2 Affinity Gel | Millipore-Sigma | Cat# A2220 |
| Microsep Advance Centrifugal Filter 10K | Pall Life Sciences | Cat# MCP010C41 |
| Superdex 200 10/300 GL HPLC Column | GE Healthcare | Cat# 17-5175-01 |

### EXPERIMENTAL MODEL AND SUBJECT DETAILS

#### Animals

All animal work was approved by and performed in accordance with the Canadian Council of Animal Care at the University of Alberta (AUP#3358). Sprague-Dawley timed pregnant rats were obtained from Charles River Laboratories and arrived at our facility one week prior to birth. Targeted replacement rats expressing human ApoE2, ApoE3 and ApoE4 were bred in house. All experiments were performed on male and female animals.

#### Primary culture of hippocampal neurons and astrocytes

Primary hippocampal cultures were prepared as previously described^1^. Briefly, hippocampi were dissected from P0-P1 rat pups and digested with papain and benzonase nuclease. After digestion, the tissues were gently triturated and filtered with a 70-µm nylon cell strainer. Neurons and glia were grown on poly-D-lysine (PDL) coated 25-mm round glass coverslips for microscopy experiments or plastic tissue culture dishes for biochemical analysis. Neurons were grown in Neurobasal medium containing B-27 supplement, 2 mM GlutaMax and antibiotic-antimycotic. 1 μM AraC was added to the media 2 days after plating and maintained for 4 days to prevent growth of glia in neuronal cultures. Astrocytes were grown in Basal Eagle Media containing 10% fetal bovine serum, 0.45% glucose, 1 mM sodium pyruvate, 2 mM GlutaMax and antibiotic-antimycotic. All cells were grown at 37°C in 5% CO_2_ and used at DIV 6-9.

#### ApoE particle characterization

ApoE particles were provided by Kisbee Therapeutics as previously described^10^, using human ApoE2, ApoE3 or ApoE4 secreted by Expi293 cells, combined with 1-palmitoyl-2-oleoyl-glycero-3-phosphocholine (POPC) at an ApoE/POPC ratio of 1:95–1:125 mol/mol. The diameter of ApoE particles was analyzed by transmission electron microscopy (TEM). Carbon grids were prepared by undergoing glow discharge in a Cressington 208 carbon coater for 40 seconds and 10 mA and 0.1 mbar. Samples were diluted in PBS to 0.05 mg/mL protein concentration and 5 μl of the sample was pipetted onto the prepared carbon grid. The grids were washed twice with H_2_O and stained with 1% uranyl acetate in H_2_O for 1 min. The grids were air dried overnight and imaged using a transmission electron microscope (JEOL JEM-2100, Gattan Orius camera with Digital micrograph) at 200 kV acceleration voltage.

#### PCSK9 purification

Wild-type human PCSK9 was purified from HEK 293S as previously described^62,85^. Briefly, HEK 293S cells stably expressing wild-type PCSK9 containing a FLAG-tag (DYKDDDDK) at the C-terminus were cultured in Dulbecco’s Modified Eagle’s Medium (DMEM) – high glucose with 10% fetal bovine serum. PSCK9 was purified by anti-FLAG M2 affinity gel chromatography according to the manufacturer’s instructions. Size-exclusion chromatography was used to isolate PCSK9 on a Tricorn Superdex 200 10/300 column. A 10 kDa-MW cut-off PALL Microsep™ filter was used to concentrate PCSK9, and protein purity was determined by SDS-PAGE and protein staining using EZ-Run protein gel staining solution.

#### Immunostaining protocol

Cells were fixed in 4% paraformaldehyde (PFA) for 10 min at room temperature, washed twice in PBS with 0.1% Triton X-100 and blocked with PBS containing 2% bovine serum albumin and 0.2% Triton X-100 for 1 h at room temperature. Cells were incubated with primary antibody diluted in blocking buffer for 1 h at room temperature, washed three times in PBS with 0.1% Triton X-100 and incubated with secondary antibody diluted in blocking buffer for 1 h at room temperature. After washing, the cells were incubated with DAPI diluted in PBS for 20 min and mounted using DAKO fluorescence mounting media. For quantification, 10 images per coverslip were averaged.

#### Confocal microscopy

Imaging was performed using the Laser Scanning Confocal Microscope (LSM) 900 with Airyscan 2 equipped with a plan-apochromat 20x air objective (Zeiss, NA = 0.8), 40x oil objective (Zeiss, NA = 1.3) and 63x oil objective (Zeiss, NA = 1.4) and ZEN software (Zeiss).

#### Cell specificity staining

Neurons and glia plated on PDL-coated glass coverslips were washed twice in PBS, fixed in 4% PFA and immunostained for β3-tubulin, GFAP or Iba1 as described above. Imaging was performed using a 20x air objective and SR mode. The imaging field of view was selected using the DAPI channel to blind the experimenter to the β3-tubulin, GFAP and Iba1 channels. 10 images were averaged per coverslip with multiple cells per image. For quantification, the precent distribution of cell type in the culture was expressed as the number of β3-tubulin-positive neurons, GFAP-positive astrocytes or Iba1-positive microglia divided by the total number of DAPI.

#### Fatty acid transfer assays

Neurons plated on PDL-coated glass coverslips were incubated with complete media containing 2 μM BODIPY 558/568 C12 (Red-C12) for 16 h, washed twice in warm PBS and incubated with fresh complete media for 1 h. Red-C12 labelled neurons and unlabelled astrocytes on separate coverslips were washed twice with warm PBS and the coverslips were sandwiched together (facing each other) separated by paraffin wax and incubated in HBSS or neuronal complete media for 4 h at 37°C with or without NaOH, 2 μM Aβ42 and/or 50 μg/ml ApoE particles. Astrocytes were fixed in 4% PFA and imaged. Imaging was performed using a 63x oil objective and SR-2Y mode. The imaging field of view was selected using the DAPI channel to blind the experimenter to the Red-C12 channel. 10 images were averaged per coverslip with 1-2 cells per image. For quantification, maximum intensity projections of three-dimensional image stacks were generated. The images were thresholded using the Yen algorithm and the number of Red-C12-positive puncta per nuclei were analyzed using Fiji-ImageJ software.

#### Lipid droplet co-localization assays

For Red-C12 co-localization, neurons plated on PDL-coated glass coverslips were incubated with complete media containing 2 μM Red-C12 for 16 h, washed twice in warm PBS and incubated with fresh complete media for 1 h. Red-C12 labelled neurons and unlabelled astrocytes on separate coverslips were washed twice with warm PBS and the coverslips were sandwiched together (facing each other) separated by paraffin wax and incubated in HBSS for 4 h at 37°C. Astrocytes were fixed in 4% PFA and stained with 5 μg/ml BODIPY 493/503 (BD493) for 1 h at room temperature. Imaging was performed using a 63x oil objective and SR-4Y mode. Quantification was performed by creating binary masks of Red-C12 and BD493 channels and generating an overlap image using the “AND” function in Fiji-ImageJ software. Percent co-localization was expressed as the number of overlapping puncta divided by the number of Red-C12 puncta. For mApple-6xHp co-localization, astrocytes plated on PDL-coated glass coverslips were transduced with 1 μg mApple-6xHp^23^ with Lipofectamine 3000 Transfection Reagent at DIV 7 according to the manufacturer’s protocol. The media was replaced with fresh media after 3 h and the cells were used 2 days later (DIV 9). Cells were fixed in 4% pre-warmed PFA and stained with 5 μg/ml BODIPY 493/503 for 1 h at room temperature. Imaging was performed using a 40x oil objective and SR-4Y mode. Quantification of co-localization between mApple-6xHp and BD493 was performed by subtracting the mApple-6xHp background using a rolling ball radius of 15 pixels and then as described above. Percent co-localization was expressed as the number of overlapping puncta divided by the number of BD493 puncta. For all co-localization assays, the imaging field of view was selected using the DAPI channel to blind the experimenter to the other channels. 10 images were averaged per coverslip with 1-2 cells per image.

#### ApoE endolysosomal co-localization assays

For GFP-LC3 co-localization, astrocytes plated on PDL-coated glass coverslips were transduced with 0.5 μg GFP-LC3^86^ with Lipofectamine 3000 Transfection Reagent at DIV 3 according to the manufacturer’s protocol. The media was replaced with fresh media after 3 h and the cells were used 3 days later (DIV 6). Cells were washed twice in warm PBS and incubated with 50 μg/ml ApoE4 particles in HBSS for 4 h at 37°C. Cells were fixed in 4% pre-warmed PFA and immunostained for ApoE as described above. Quantification was performed by creating binary masks of GFP-LC3 and ApoE channels and generating an overlap image using the “AND” function in Fiji-ImageJ software. Percent co-localization was expressed as the number of overlapping puncta divided by the number of GFP-LC3 puncta. For LAMP1, Rab5 or Rab7 co-localization, astrocytes plated on PDL-coated glass coverslips were washed twice in warm PBS incubated with 50 μg/ml ApoE2 or ApoE4 particles in HBSS for 4 h at 37°C. Cells were fixed in 4% PFA and immunostained for ApoE and either LAMP1, Rab5 or Rab7 as described above. Quantification was performed by creating binary masks of ApoE and either LAMP1, Rab5 or Rab7 channels and generating an overlap image using the “AND” function in Fiji-ImageJ software. Percent co-localization was expressed as the number of overlapping puncta divided by the number of ApoE puncta. For all co-localization assays, imaging was performed using a 63x oil objective and SR-2Y mode. The imaging field of view was selected using the DAPI channel to blind the experimenter to the other channels. 10 images were averaged per coverslip with 1-2 cells per image.

#### Autofluorescence lipofuscin co-localization assays

Astrocytes plated on PDL-coated glass coverslips were washed twice in warm PBS incubated with 50 μg/ml ApoE4 particles in HBSS for 4 h at 37°C. Cells were fixed in 4% PFA and immunostained for Rab7 as described above. Imaging was performed using a 63x oil objective and SR-2Y mode. Autofluorescence was detected using 488 nm excitation. The imaging field of view was selected using the DAPI channel to blind the experimenter to the other channels. 10 images were averaged per coverslip with 1-2 cells per image. Quantification was performed by creating binary masks of autofluorescent lipofuscin and Rab7 channels and generating an overlap image using the “AND” function in Fiji-ImageJ software. Percent co-localization was expressed as the number of overlapping puncta divided by the number of lipofuscin puncta.

#### Lipofuscin transmission electron microscopy

Astrocytes plated on 1 x 1 PDL-coated aclar sheets were washed twice in warm PBS and incubated with 50 μg/ml ApoE4 particles in HBSS for 4 h at 37°C. Cells were washed twice in warm PBS and fixed in 1% PFA, 2% glutaraldehyde in 0.1 M CaB (0.1–0.2 M Cacodylate buffer, pH 7.4) containing 4% polyvinylpyrrolidone and 0.05% CaCl_2_ for 16 h at 4°C. Cells were washed with 0.1 M CaB and post-fixed with 2% osmium-imidazole buffer, pH 7.5. Samples were dehydrated in a graded series of ethanol, 30%, 50%, and then block stained with 1% uranyl acetate in 70% ethanol overnight. Samples were then further dehydrated in 70, 95, and 100% ethanol and anhydrous acetone and embedded in Durcupan (ACM) resin. Blocks were sectioned using an ultramicrotome (Leica, EM UC6; 70 nm thickness) and placed on a 400 mesh copper grid. Sections were stained with 1% uranyl acetate and 1% lead citrate and imaged using a transmission electron microscope (JEOL JEM-2100, Gatan Orius camera with Digital micrograph) at 200 kV acceleration voltage.

#### Exogenous fatty acid release assays

Astrocytes were grown on PDL-coated 6-well plastic dishes. Astrocytes were loaded with 2 μM Red-C12 for 16 h in complete media. Cells were washed twice in warm PBS, incubated with fresh media for 1 h and treated with or without 50 μg/ml ApoE particles in HBSS for 4 h at 37°C. Astrocyte-conditioned media was collected and centrifuged at 16,000 x g for 15 min to remove dead cells or cell debris. The supernatant was analyzed for Red-C12 (excitation/emission 561/600 nm) fluorescence using Synergy Mx Multi-Mode Microplate Reader (BioTek Instruments Inc.). Each treatment was analyzed in triplicate and averaged for each biological replicate.

#### Lipid droplet assays

Astrocytes plated on PDL-coated glass coverslips were washed twice in warm PBS and treated with or without DMSO, 50 μg/ml ApoE particles, 100 nM baf A1, 10 μM rimeporide or 200 nM YM-201636 in HBSS for 4 h at 37°C. For the PCSK9 lipid droplet assays, astrocytes plated on PDL-coated glass coverslips, were washed twice in warm PBS and pre-treated with or without 10 μg/ml PCSK9 in HBSS for 2 h at 37°C. Astrocytes were then treated with or without 50 μg/ml ApoE4 particles or 10 μg/ml PCSK9 in HBSS for 4 h at 37°C. Cells were fixed in 4% PFA and stained with 5 μg/ml BODIPY 493/503 for 1 h at room temperature. Imaging was performed using a 63x oil objective and SR-2Y mode. The imaging field of view was selected using the DAPI channel to blind the experimenter to the other channels. 10 images were averaged per coverslip with 1-2 cells per image. The ApoE particle reduced serum lipid droplet assays were imaged and quantified blind. The PCSK9 and YM-201636 lipid droplet assays were quantified blind. For quantification, maximum intensity projections of three-dimensional image stacks were generated. The images were thresholded using the MaxEntropy algorithm and the number of BD493-positive lipid droplets per nuclei and average size of lipid droplets were analyzed using Fiji-ImageJ software.

#### LysoTracker acidification assay

Astrocytes plated on PDL-coated glass coverslips were washed twice in warm PBS and treated with or without DMSO, 100 nM bafilomycin A1 (baf A1), 200 nM bafilomycin A1, 10 μM rimeporide, 200 nM YM-201636 or 50 μg/ml ApoE particles in HBSS for 4 h at 37°C with 50 nM LysoTracker Red DND-99 added for the last 30 min. Cells were fixed in 4% PFA. Imaging was performed using a 63x oil objective and SR-4Y mode. The imaging field of view was selected using the DAPI channel to blind the experimenter to the LysoTracker Red channel. 10 images were averaged per coverslip with 1-2 cells per image. For quantification, maximum intensity projections of three-dimensional image stacks were generated. The images were thresholded using the MaxEntropy algorithm and the number of LysoTracker Red-positive puncta per nuclei and average size of puncta were analyzed using Fiji-ImageJ software.

#### Cathepsin D assay

Astrocytes were grown on PDL-coated 6-well plastic dishes. Astrocytes were washed twice in warm PBS and incubated in HBSS with or without DMSO, 10 μM rimeporide, 200 nM YM-201636 or 50 μg/ml ApoE particles for 4 h at 37°C. Lysates were analyzed using a cathepsin D activity assay kit according to the manufacturer’s protocol and Synergy Mx Multi-Mode Microplate Reader (BioTek Instruments Inc.). Each treatment was analyzed in duplicate and averaged for each biological replicate. The assays were normalized to the control treatment for each biological replicate.

#### Triglyceride assay

Astrocytes were grown on PDL-coated 6-well plastic dishes. Astrocytes were washed twice in warm PBS and incubated in HBSS with or without DMSO, 50 μg/ml ApoE particles, 10 μM rimeporide or 200 nM YM-201636 in HBSS for 4 h at 37°C. Lysates were collected and total triglyceride content was analyzed using a triglyceride-glo assay kit according to the manufacturer’s protocol and Synergy Mx Multi-Mode Microplate Reader (BioTek Instruments Inc.). Each treatment was analyzed in duplicate and averaged for each biological replicate. The assays were normalized to lysate protein levels as calculated by a BCA assay.

#### Autofluorescence lipofuscin assays

For the ApoE particle, bafilomycin A1 (baf A1), rimeporide and YM-201636 assays, astrocytes plated on PDL-coated glass coverslips were washed twice in warm PBS and treated with or without DMSO, 50 μg/ml ApoE particles, 100 nM baf A1, 10 μM rimeporide or 200 nM YM-201636 in HBSS for 4 h at 37°C. For the PCSK9 assays, astrocytes plated on PDL-coated glass coverslips, were washed twice in warm PBS and pre-treated with or without 10 μg/ml PCSK9 in HBSS for 2 h at 37°C. Astrocytes were then treated with or without 50 μg/ml ApoE4 particles or 10 μg/ml PCSK9 in HBSS for 4 h at 37°C. Cells were fixed in 4% PFA. Imaging was performed using a 40x oil objective and SR-4Y mode. Autofluorescence was detected using 488 nm excitation. The imaging field of view was selected using the DAPI channel to blind the experimenter to the autofluorescent channel. 10 images were averaged per coverslip with multiple cells per image. All images were quantified blind. For quantification, sum slice projections of three-dimensional image stacks were generated. The images were manually thresholded and the number and area of autofluorescent lipofuscin puncta were analyzed using Fiji-ImageJ software.

#### ApoE particle internalization assay

Astrocytes were grown on PDL-coated 6-well plastic dishes. Astrocytes were washed twice in warm PBS and treated with or without 25 μg/ml ApoE particles in HBSS for 0-, 15- or 30-min. Glia were washed twice in ice-cold acid wash (0.5 M NaCl and 0.2 M acetic acid pH 2.5), followed by two washes with ice-cold PBS. Astrocytes were lysed in ice-cold lysis buffer (20 mM HEPES pH 7.4, 100 mM NaCl, 1% Triton X-100, 5 mM EDTA, HALT Protease Inhibitor), resolved by SDS-PAGE and processed for western blotting using anti-ApoE goat polyclonal and anti-β-actin mouse monoclonal as a loading control.

#### PCSK9 ApoE particle internalization assay

Astrocytes were grown on PDL-coated 6-well plastic dishes. Astrocytes were washed twice in warm PBS and treated with or without 10 μg/ml PCSK9 in HBSS for 4 h at 37°C with 50 μg/ml ApoE4 particles added for the last 15 or 0 min. Astrocytes were washed twice in ice-cold acid wash (0.5 M NaCl and 0.2 M acetic acid pH 2.5), followed by two washes with ice-cold PBS. Astrocytes were lysed in ice-cold lysis buffer (20 mM HEPES pH 7.4, 100 mM NaCl, 1% Triton X-100, 5 mM EDTA, HALT Protease Inhibitor), resolved by SDS-PAGE and processed for western blotting using anti-ApoE goat polyclonal and anti-β-actin mouse monoclonal as a loading control.

#### Statistics

Datasets were assembled in Microsoft Excel 365 (Microsoft Corp.). Statistical analysis and graphing were performed using Graphpad Prism 10. All graphs are depicted as Superplots where the independent replicates are shown in large shapes, and the corresponding technical replicants shown as small shapes of the same color ^87^. The independent replicate values were calculated from the mean of technical replicates within an experiment. Statistical analyses were performed on the independent replicates. Statistical test used and p-values and non-significant comparisons can be found in the figures and/or figure legends. Where stated the Bonferroni correction method was used to correct for multiple comparisons (raw p-values were multiplied by the number of comparisons in the experiment).

---

**Figure S1.**
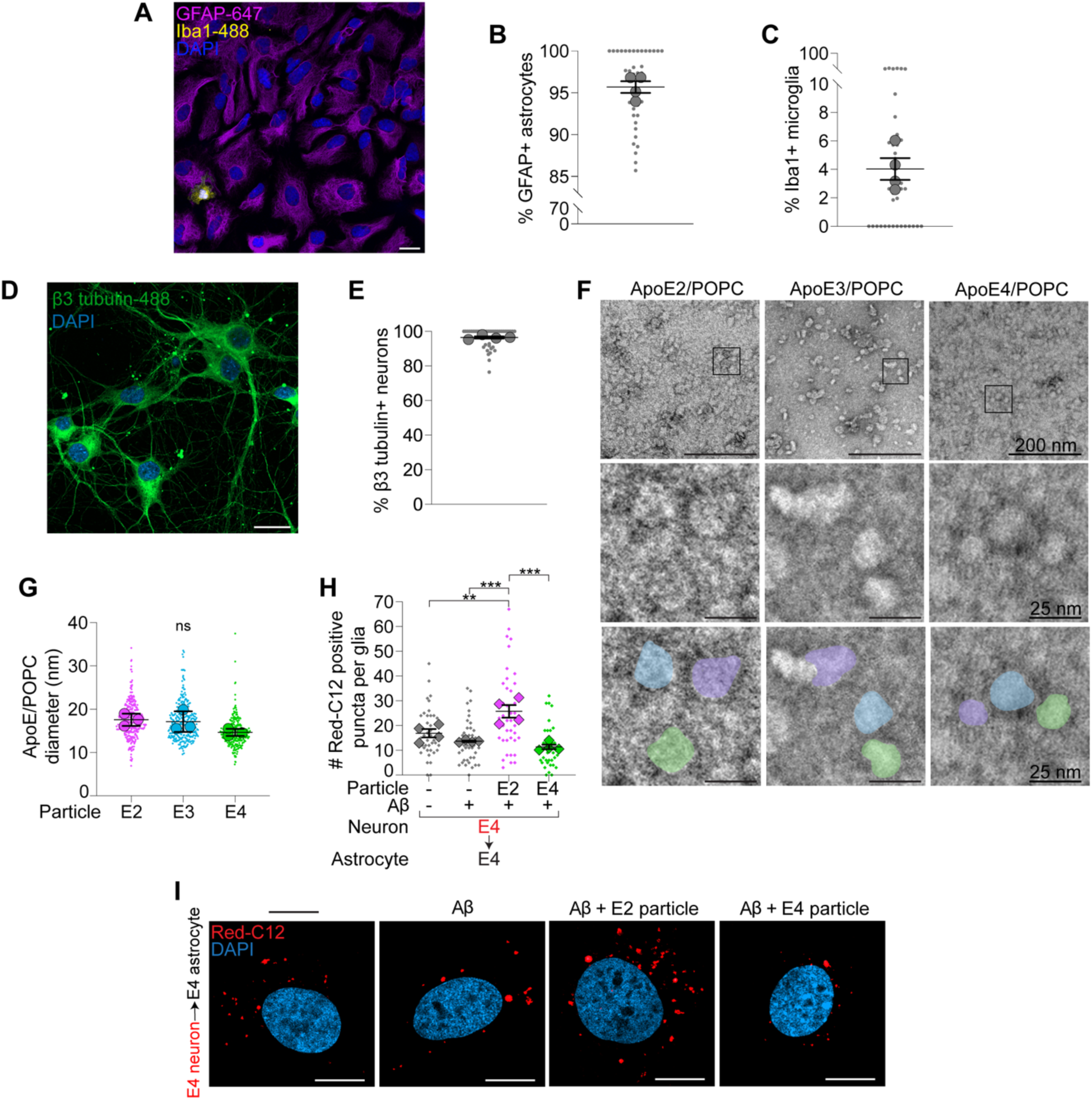
Validation of ApoE particles and primary astrocyte cultures. (A) Representative airyscan image of glial culture immunostained for GFAP and Iba1. Scale bar is 20 μm. (B and C) Percentage of DAPI-positive cells also positive for GFAP astrocytes or Iba1 microglia in cultured glia. n = 4 independent replicates; mean ± SEM. (D) Representative airyscan image of neuronal culture immunostained for neuron-specific β3-tubulin. Scale bar is 20 μm. (E) Percentage of DAPI-positive cells also positive for β3-tubulin in cultured neurons. n = 4 independent replicates; mean ± SEM. (F) ApoE particles imaged by transmission electron microscopy (TEM). Boxed areas magnified in bottom panels. Shaded regions highlight particles. Scale bars are 200 nm and 25 nm. (G) Size distribution of ApoE particle diameters determined by TEM. n = 3 technical replicates; mean ± SD. One-way ANOVA with Tukey’s post-test. ns, no significant differences were discovered. (H) Red-C12 positive puncta in ApoE4 astrocytes following Red-C12 transfer assay with ApoE4 neurons in media ± Aβ or 50 μg/ml ApoE2 or ApoE4 particles. n = 4 independent experiments; mean ± SEM. One-way ANOVA with Tukey’s post-test. (I) Airyscan images of ApoE4 astrocytes displayed as maximum intensity projections following Red-C12 transfer assay in media ± Aβ or 50 μg/ml ApoE2 or ApoE4 particles. Scale bars are 10 μm. For all graphs independent replicates are in large shapes and technical replicates in small shapes; *p < 0.05, **p < 0.01, ***p < 0.001, ****p < 0.0001.

**Figure S2.**
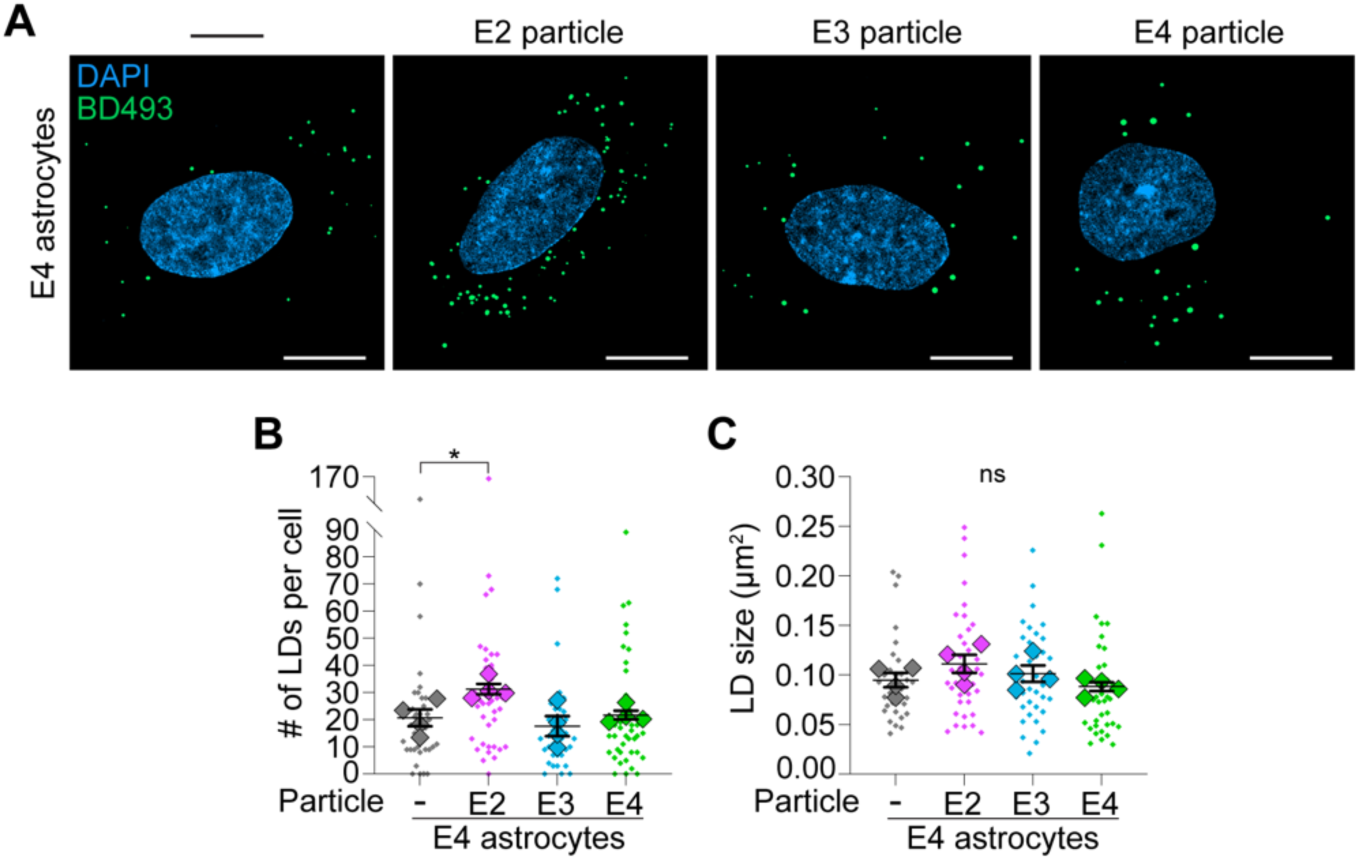
No ApoE-isoform specific differences on lipid storage in ApoE4 astrocytes. (A-C) Airyscan images of ApoE4 astrocytes ± 50 μg/ml ApoE2, ApoE3 or ApoE4 particle treatment in HBSS stained for BD493-positive lipid droplets (LDs). n = 4 independent experiments; mean ± SEM. One-way ANOVA with Dunnett’s post-test. Scale bars are 10 μm. All images displayed as maximum intensity projections. For all graphs independent replicates are in large shapes and technical replicates in small shapes; *p < 0.05. ns, no significant differences were discovered.

**Figure S3.**
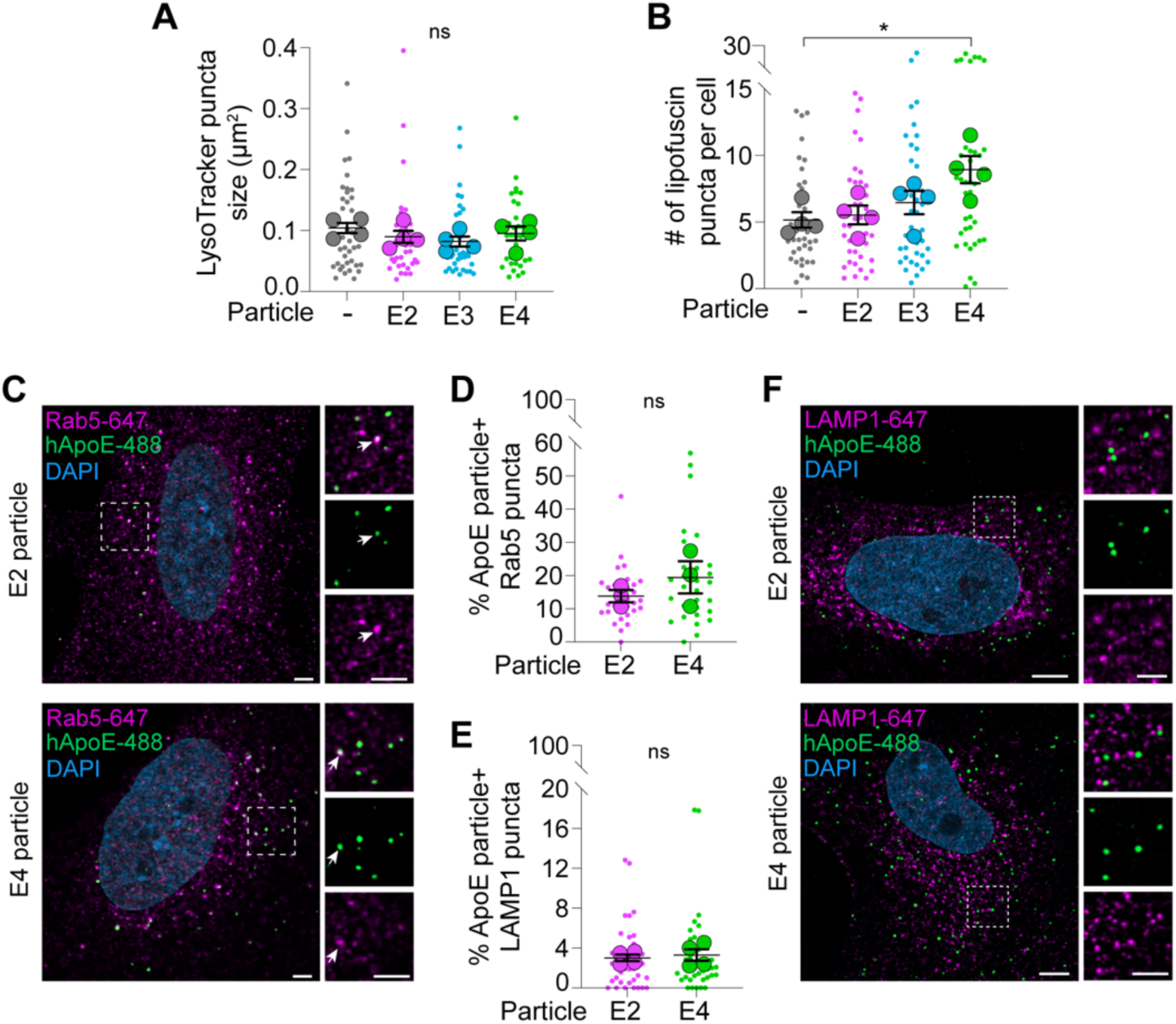
ApoE4 particles impair endolysosomal function. (A) Quantification of LysoTracker Red-positive puncta size in astrocytes ± 50 μg/ml ApoE2, ApoE3 or ApoE4 particles in HBSS. n = 4 independent experiments; mean ± SEM. One-way ANOVA with Tukey’s post-test. (B) Quantification of autofluorescent lipofuscin puncta number in astrocytes ± 50 μg/ml ApoE2, ApoE3 or ApoE4 particles in HBSS. n = 4 independent experiments; mean ± SEM. One-way ANOVA with Tukey’s post-test. (C and D) Airyscan image of astrocyte ± 50 μg/ml ApoE2 or ApoE4 particles in HBSS showing ApoE and Rab5. Boxed area magnified in right panels. Arrows highlight co-localization. The percent co-localization of human ApoE (hApoE) and Rab5 puncta were quantified. n = 3 independent replicates; mean ± SEM. Two-tailed Student’s t test. Scale bars are 2.5 μM. (E) Percent co-localization of ApoE and LAMP1 puncta in astrocytes ± 50 μg/ml ApoE2 or ApoE4 particles in HBSS. n=4 independent experiments; mean ± SEM. Two-tailed Student’s t test (F) Airyscan image of astrocyte ± 50 μg/ml ApoE2 or ApoE4 particles in HBSS showing human ApoE (hApoE) and LAMP1. Boxed area magnified in right panels. Scale bars are 5 μM. For all graphs independent replicates are in large shapes and technical replicates in small shapes; *p < 0.05, **p < 0.01, ***p < 0.001, ****p < 0.0001. ns, no significant differences were discovered.

**Figure S4.**
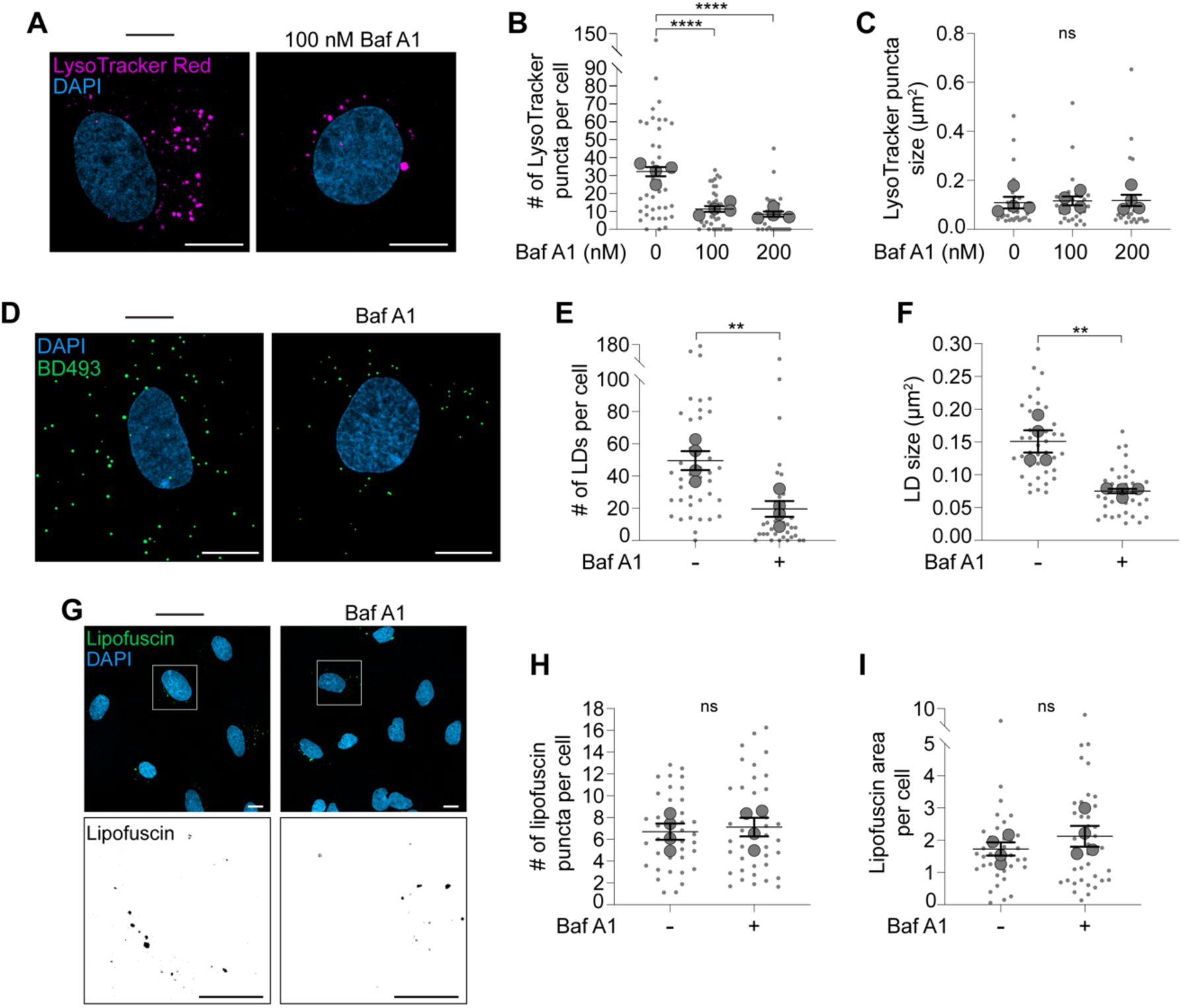
Impairing endolysosomal function mimics the effects of ApoE4 particles. (A-C) Airyscan images of astrocytes ± bafilomycin A1 (baf A1) in HBSS stained for LysoTracker Red. n = 4 independent experiments: mean ± SEM. One-way ANOVA with Tukey’s post-test. Scale bars are 10 μm. (D-F) Airyscan images of astrocytes ± baf A1 in HBSS stained for BD493-positive lipid droplets (LDs). n = 4 independent replicates; mean ± SEM. Two-tailed Student’s t test. Scale bars are 10 μm. (G-I) Airyscan images of autofluorescent lipofuscin in astrocytes ± baf A1 in HBSS. Boxed area magnified in bottom panels. n = 4 independent experiments; mean ± SEM. Two-tailed Student’s t test. Scale bars are 10 μm. All images displayed as maximum intensity projections. For all graphs independent replicates are in large shapes and technical replicates in small shapes; *p < 0.05, **p < 0.01, ***p < 0.001, ****p < 0.0001. ns, no significant differences were discovered.

**Figure S5.**
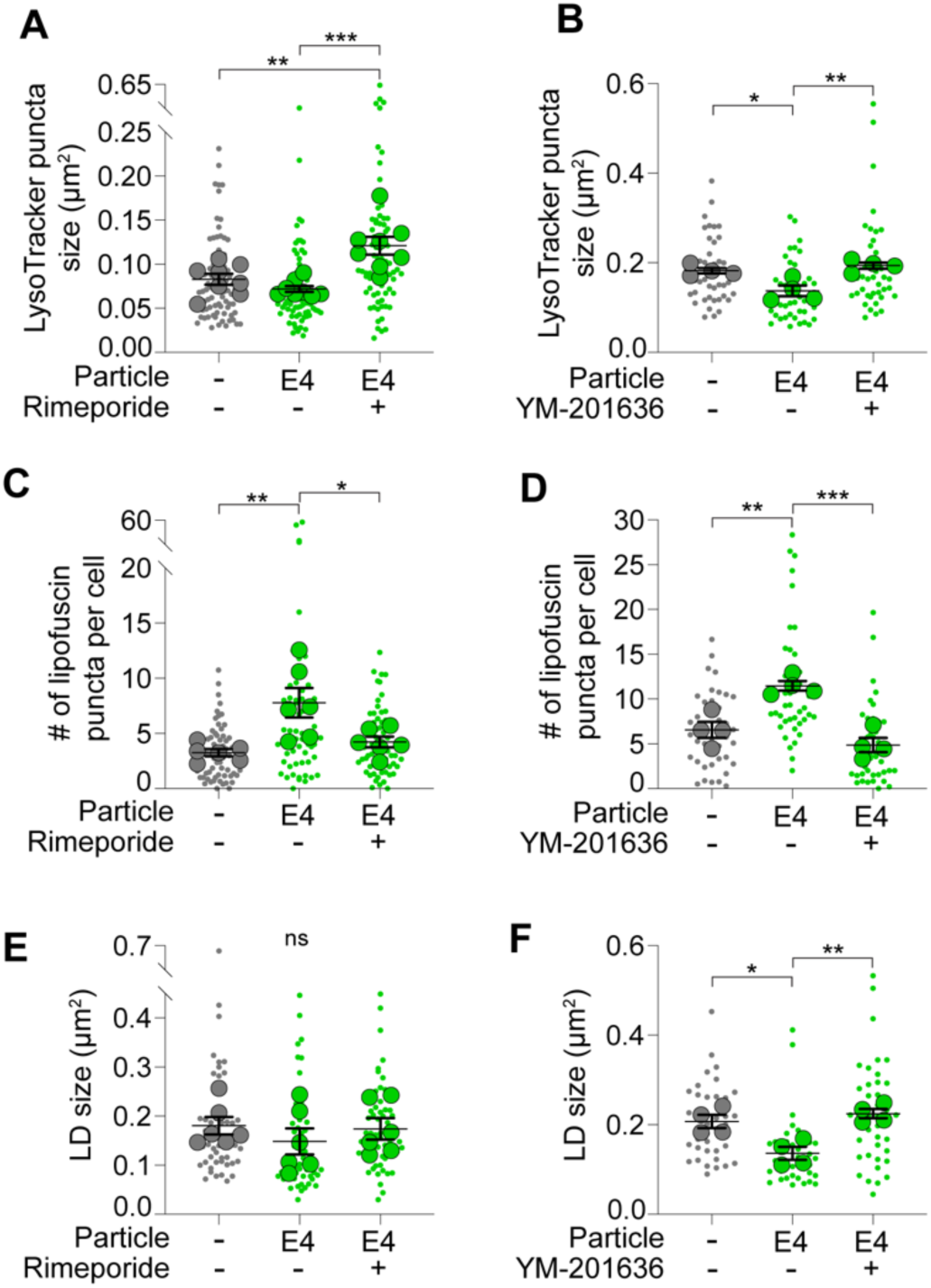
Restoring endosomal function prevents ApoE4-mediated alterations. (A) Quantification of LysoTracker Red puncta size in astrocytes ± rimeporide or 50 μg/ml ApoE4 particles in HBSS. n = 8 independent experiments; mean ± SEM. One-way ANOVA with Tukey’s post-test. (B) Quantification of LysoTracker Red puncta size in astrocytes ± YM-201636 or 50 μg/ml ApoE4 particles in HBSS. n = 4 independent experiments; mean ± SEM. One-way ANOVA with Tukey’s post-test. (C) Quantification of autofluorescent lipofuscin puncta number in astrocytes ± rimeporide or 50 μg/ml ApoE4 particles in HBSS. n = 6 independent experiments; mean ± SEM. One-way ANOVA with Tukey’s post-test. (D) Quantification of autofluorescent lipofuscin puncta number in astrocytes ± YM-201636 or 50 μg/ml ApoE4 particles in HBSS. n = 4 independent experiments; mean ± SEM. One-way ANOVA with Tukey’s post-test. (E) Quantification of BD493-positive lipid droplet (LD) size in astrocytes ± rimeporide or 50 μg/ml ApoE4 particle treatment in HBSS. n = 6 independent experiments; mean ± SEM. One-way ANOVA with Tukey’s post-test. (F) Quantification of BD493-positive lipid droplet (LD) size in astrocytes ± YM-201636 or 50 μg/ml ApoE4 particle treatment in HBSS. n = 4 independent experiments; mean ± SEM. One-way ANOVA with Tukey’s post-test. For all graphs independent replicates are in large shapes and technical replicates in small shapes; *p < 0.05, **p < 0.01, ***p < 0.001, ****p < 0.0001. ns, no significant differences were discovered.

## REFERENCES

1. Ioannou, M.S., Jackson, J., Sheu, S.-H., Chang, C.-L., Weigel, A. V, Liu, H., Pasolli, H.A., Xu, C.S., Pang, S., Matthies, D., et al. (2019). Neuron-Astrocyte Metabolic Coupling Protects against Activity-Induced Fatty Acid Toxicity. Cell 177, 1522–1535.e14. 10.1016/j.cell.2019.04.001.

2. Liu, L., MacKenzie, K.R., Putluri, N., Maletić-Savatić, M., and Bellen, H.J. (2017). The Glia-Neuron Lactate Shuttle and Elevated ROS Promote Lipid Synthesis in Neurons and Lipid Droplet Accumulation in Glia via APOE/D. Cell Metab. 26, 719–737.e6. 10.1016/j.cmet.2017.08.024.

3. Van Den Brink, D.M., Cubizolle, A., Chatelain, G., Davoust, N., Girard, V., Johansen, S., Napoletano, F., Dourlen, P., Guillou, L., Angebault-Prouteau, C., et al. (2018). Physiological and pathological roles of FATP-mediated lipid droplets in Drosophila and mice retina. PLoS Genet. 14. 10.1371/journal.pgen.1007627.

4. Pitas, R.E., Boyles, J.K., Lee, S.H., Foss, D., and Mahley, R.W. (1987). Astrocytes synthesize apolipoprotein E and metabolize apolipoprotein E-containing lipoproteins. Biochimica et Biophysics Actcr 917, 148–161.

5. Flowers, S.A., and Rebeck, G.W. (2020). APOE in the normal brain. Preprint at Academic Press Inc., 10.1016/j.nbd.2019.104724 10.1016/j.nbd.2019.104724.

6. Windham, I.A., and Cohen, S. (2024). The cell biology of APOE in the brain. Preprint at Elsevier Ltd, 10.1016/j.tcb.2023.09.004 10.1016/j.tcb.2023.09.004.

7. Xu, Q., Bernardo, A., Walker, D., Kanegawa, T., Mahley, R.W., and Huang, Y. (2006). Profile and regulation of apolipoprotein E (ApoE) expression in the CNS in mice with targeting of green fluorescent protein gene to the ApoE locus. The Journal of Neuroscience 26, 4985–4994. 10.1523/JNEUROSCI.5476-05.2006.

8. Qi, G., Mi, Y., Shi, X., Gu, H., Brinton, R.D., and Yin, F. (2021). ApoE4 Impairs Neuron-Astrocyte Coupling of Fatty Acid Metabolism. Cell Rep. 34. 10.1016/j.celrep.2020.108572.

9. Chen, H., Zhao, S., Jian, Q., Yan, Y., Wang, S., Zhang, X., and Ji, Y. (2024). The role of ApoE in fatty acid transport from neurons to astrocytes under ischemia/hypoxia conditions. Mol. Biol. Rep. 51. 10.1007/s11033-023-08921-4.

10. Ralhan, I., Do, A.D., Bae, J.-Y., Feringa, F.M., Cai, W., Chang, J., Chik, K., Lee, N.Y.J., Gerry, C.J., van der Kant, R., et al. (2025). Protective ApoE variants support neuronal function by effluxing oxidized phospholipids. Neuron. 10.1016/j.neuron.2025.10.040.

11. Herz, J., and Bock, H.H. (2002). Lipoprotein receptors in the nervous system. Preprint, 10.1146/annurev.biochem.71.110601.135342 10.1146/annurev.biochem.71.110601.135342.

12. Rambold, A.S., Cohen, S., and Lippincott-Schwartz, J. (2015). Fatty acid trafficking in starved cells: Regulation by lipid droplet lipolysis, autophagy, and mitochondrial fusion dynamics. Dev. Cell 32, 678–692. 10.1016/j.devcel.2015.01.029.

13. Fryer, J.D., DeMattos, R.B., McCormick, L.M., O’Dell, M.A., Spinner, M.L., Bales, K.R., Paul, S.M., Sullivan, P.M., Parsadanian, M., Bu, G., et al. (2005). The low density lipoprotein receptor regulates the level of central nervous system human and murine apolipoprotein E but does not modify amyloid plaque pathology in PDAPP mice. Journal of Biological Chemistry 280, 25754–25759. 10.1074/jbc.M502143200.

14. Guo, J.L., Braun, D., Fitzgerald, G.A., Hsieh, Y.T., Rougé, L., Litvinchuk, A., Steffek, M., Propson, N.E., Heffner, C.M., Discenza, C., et al. (2025). Decreased lipidated ApoE-receptor interactions confer protection against pathogenicity of ApoE and its lipid cargoes in lysosomes. Cell 188, 187–206.e26. 10.1016/j.cell.2024.10.027.

15. Nuriel, T., Peng, K.Y., Ashok, A., Dillman, A.A., Figueroa, H.Y., Apuzzo, J., Ambat, J., Levy, E., Cookson, M.R., Mathews, P.M., et al. (2017). The endosomal-lysosomal pathway is dysregulated by APOE4 expression in vivo. Front. Neurosci. 11. 10.3389/fnins.2017.00702.

16. Grefhorst, A., McNutt, M.C., Lagace, T.A., and Horton, J.D. (2008). Plasma PCSK9 preferentially reduces liver LDL receptors in mice. J. Lipid Res. 49, 1303–1311. 10.1194/jlr.M800027-JLR200.

17. Jaafar, A.K., Techer, R., Chemello, K., Lambert, G., and Bourane, S. (2023). PCSK9 and the nervous system: a no-brainer? Preprint at American Society for Biochemistry and Molecular Biology Inc., 10.1016/j.jlr.2023.100426 10.1016/j.jlr.2023.100426.

18. Quinlivan, V.H., Wilson, M.H., Ruzicka, J., and Farber, S.A. (2017). An HPLC-CAD/fluorescence lipidomics platform using fluorescent fatty acids as metabolic tracers. J. Lipid Res. 58, 1008–1020. 10.1194/jlr.D072918.

19. Ioannou, M.S., Liu, Z., and Lippincott-Schwartz, J. (2019). A Neuron-Glia Co-culture System for Studying Intercellular Lipid Transport. Curr. Protoc. Cell Biol. 84, e95. 10.1002/cpcb.95.

20. Strickland, M.R., Rau, M.J., Summers, B., Basore, K., Wulf, J., Jiang, H., Chen, Y., Ulrich, J.D., Randolph, G.J., Zhang, R., et al. (2024). Apolipoprotein E secreted by astrocytes forms antiparallel dimers in discoidal lipoproteins. Neuron 112, 1100–1109.e5. 10.1016/j.neuron.2023.12.018.

21. Hampel, H., Hardy, J., Blennow, K., Chen, C., Perry, G., Kim, S.H., Villemagne, V.L., Aisen, P., Vendruscolo, M., Iwatsubo, T., et al. (2021). The Amyloid-β Pathway in Alzheimer’s Disease. Mol. Psychiatry 26, 5481–5503. 10.1038/s41380-021-01249-0.

22. Kuperstein, I., Broersen, K., Benilova, I., Rozenski, J., Jonckheere, W., Debulpaep, M., Vandersteen, A., Segers-Nolten, I., Van Der Werf, K., Subramaniam, V., et al. (2010). Neurotoxicity of Alzheimer’s disease Aβ peptides is induced by small changes in the Aβ42 to Aβ40 ratio. EMBO Journal 29, 3408–3420. 10.1038/emboj.2010.211.

23. Li, X., Gamuyao, R., Wu, M.L., Cho, W.J., King, S. V., Petersen, R.A., Stabley, D.R., Lindow, C., Climer, L.K., Shirinifard, A., et al. (2024). A fluorogenic complementation tool kit for interrogating lipid droplet-organelle interaction. J. Cell Biol. 223. 10.1083/jcb.202311126.

24. Olzmann, J.A., and Carvalho, P. (2019). Dynamics and functions of lipid droplets. Nat. Rev. Mol. Cell Biol. 20, 137–155. 10.1038/s41580-018-0085-z.

25. Walther, T.C., and Farese, R.V.J. (2012). Lipid droplets and cellular lipid metabolism. Annu. Rev. Biochem. 81, 687–714. 10.1146/annurev-biochem-061009-102430.

26. Ralhan, I., Chang, C.L., Lippincott-Schwartz, J., and Ioannou, M.S. (2021). Lipid droplets in the nervous system. Journal of Cell Biology 220. 10.1083/jcb.202102136.

27. Rubio-Atonal, L.F., Chang, J., Jacquemyn, J., Ralhan, I., Ilarraza, I., and Ioannou, M.S. (2025). Glutamate decreases oxidative stress and lipid droplet formation in astrocytes. J. Cell Sci. 138. 10.1242/jcs.263983.

28. Seehafer, S.S., and Pearce, D.A. (2006). You say lipofuscin, we say ceroid: Defining autofluorescent storage material. Preprint, 10.1016/j.neurobiolaging.2005.12.006

29. Birgisdottir, Å.B., and Johansen, T. (2020). Autophagy and endocytosis – interconnections and interdependencies. Preprint at Company of Biologists Ltd, 10.1242/jcs.228114 10.1242/jcs.228114.

30. Simonovitch, S., Schmukler, E., Bespalko, A., Iram, T., Frenkel, D., Holtzman, D.M., Masliah, E., Michaelson, D.M., and Pinkas-Kramarski, R. (2016). Impaired Autophagy in APOE4 Astrocytes. Journal of Alzheimer’s Disease 51, 915–927. 10.3233/JAD-151101.

31. Zhang, D.W., Lagace, T.A., Garuti, R., Zhao, Z., McDonald, M., Horton, J.D., Cohen, J.C., and Hobbs, H.H. (2007). Binding of proprotein convertase subtilisin/kexin type 9 to epidermal growth factor-like repeat A of low density lipoprotein receptor decreases receptor recycling and increases degradation. Journal of Biological Chemistry 282, 18602–18612. 10.1074/jbc.M702027200.

32. Zhang, D.-W., Garuti, R., Tang, W.-J., Cohen, J.C., and Hobbs, H.H. (2008). Structural requirements for PCSK9-mediated degradation of the low-density lipoprotein receptor.

33. Li, R., Zhang, C.X., Liu, J., and Zhang, D. (2025). PCSK9-promoted LDLR degradation: Recruitment or prevention of essential cofactors? Metabol. Open 26, 100362. 10.1016/j.metop.2025.100362.

34. Mauvezin, C., and Neufeld, T.P. (2015). Bafilomycin A1 disrupts autophagic flux by inhibiting both V-ATPase-dependent acidification and Ca-P60A/SERCA-dependent autophagosome-lysosome fusion. Autophagy 11, 1437–1438. 10.1080/15548627.2015.1066957.

35. Zou, J., Mitra, K., Anees, P., Oettinger, D., Ramirez, J.R., Veetil, A.T., Gupta, P.D., Rao, R., Smith, J.J., Kratsios, P., et al. (2024). A DNA nanodevice for mapping sodium at single-organelle resolution. Nat. Biotechnol. 42, 1075–1083. 10.1038/s41587-023-01950-1.

36. Samaddar, M., Fitzgerald, G.A., Nguyen, A.H., Davis, S.S., Jain, S., Guo, J., Propson, N.E., van Lengerich, B., Shi, Y., Balasundar, S., et al. (2025). Lysosomal polyamine storage upon ATP13A2 loss impairs β-glucocerebrosidase via altered lysosomal pH and electrostatic hydrolase-lipid interactions. Cell Rep. 44. 10.1016/j.celrep.2025.116179.

37. Pethő, Z., Najder, K., Beel, S., Fels, B., Neumann, I., Schimmelpfennig, S., Sargin, S., Wolters, M., Grantins, K., Wardelmann, E., et al. (2023). Acid-base homeostasis orchestrated by NHE1 defines the pancreatic stellate cell phenotype in pancreatic cancer. 10.1172/jci.

38. Tejwani, L., Balak, C., Skuja, L.L., Tatarakis, D., Rana, A., Fitzgerald, G.A., Ha, C., Lunkes de Melo, G., Sun, E.W., Heffner, C.M., et al. (2024). Lysosomes cell autonomously regulate myeloid cell states and immune responses. bioRxiv. 10.1101/2024.11.11.623074.

39. Pohlkamp, T., Xian, X., Wong, C.H., Durakoglugil, M.S., Werthmann, G.C., Saido, T.C., Evers, B.M., White, C.L., Connor, J., Hammer, R.E., et al. (2021). NHE6 depletion corrects ApoE4-mediated synaptic impairments and reduces Amyloid plaque load. Elife 10. 10.7554/eLife.72034.

40. Jacquemyn, J., Marriott, B., Chang, J., Iftikhar, E., Chik, K., Lee, N.Y.J., Rubio Atonal, L.F., Green, C., Wong, J., Acevedo-Morantes, C., et al. (2026). Glucosylceramide-induced ectosomes propagate pathogenic α-synuclein in Parkinson’s disease. Nat. Cell Biol. 10.1038/s41556-026-01871-6.

41. Wartosch, L., Fuhrmann, J.C., Schweizer, M., Stauber, T., and Jentsch, T.J. (2009). Lysosomal degradation of endocytosed proteins depends on the chloride transport protein ClC-7. FASEB Journal 23, 4056–4068. 10.1096/fj.09-130880.

42. Zhang, S., Liu, Y., Zhang, B., Zhou, J., Li, T., Liu, Z., Li, Y., and Yang, M. (2020). Molecular insights into the human CLC-7/Ostm1 transporter.

43. Wu, J.Z., Zeziulia, M., Kwon, W., Jentsch, T.J., Grinstein, S., and Freeman, S.A. (2023). ClC-7 drives intraphagosomal chloride accumulation to support hydrolase activity and phagosome resolution. Journal of Cell Biology 222. 10.1083/jcb.202208155.

44. Wu, J.Z., Pemberton, J.G., Morioka, S., Sasaki, J., Bablani, P., Sasaki, T., Balla, T., Grinstein, S., and Freeman, S.A. (2025). Sorting nexin 10 regulates lysosomal ionic homeostasis via ClC-7 by controlling PI(3,5)P2. J. Cell Biol. 224. 10.1083/jcb.202408174.

45. Feng, X., Liu, S., and Xu, H. (2023). Not just protons: Chloride also activates lysosomal acidic hydrolases. Journal of Cell Biology 222. 10.1083/jcb.202305007.

46. Zhang, Q., Li, Y., Jian, Y., Li, M., and Wang, X. (2023). Lysosomal chloride transporter CLH-6 protects lysosome membrane integrity via cathepsin activation. Journal of Cell Biology 222. 10.1083/jcb.202210063.

47. Jefferies, H.B.J., Cooke, F.T., Jat, P., Boucheron, C., Koizumi, T., Hayakawa, M., Kaizawa, H., Ohishi, T., Workman, P., Waterfield, M.D., et al. (2008). A selective PIKfyve inhibitor blocks PtdIns(3,5)P2 production and disrupts endomembrane transport and retroviral budding. EMBO Rep. 9, 164–170. 10.1038/sj.embor.7401155.

48. Leray, X., Hilton, J.K., Nwangwu, K., Becerril, A., Mikusevic, V., Fitzgerald, G., Amin, A., Weston, M.R., and Mindell, J.A. (2022). Tonic inhibition of the chloride/proton antiporter ClC-7 by PI(3,5)P2 is crucial for lysosomal pH maintenance. Elife 11. 10.7554/eLife.74136.

49. Sienski, G., Narayan, P., Bonner, J.M., Kory, N., Boland, S., Arczewska, A.A., Ralvenius, W.T., Akay, L., Lockshin, E., He, L., et al. (2021). APOE4 disrupts intracellular lipid homeostasis in human iPSC-derived glia. Sci. Transl. Med. 13. 10.1126/scitranslmed.aaz4564.

50. Windham, I.A., Powers, A.E., Ragusa, J. V., Wallace, E.D., Zanellati, M.C., Williams, V.H., Wagner, C.H., White, K.K., and Cohen, S. (2024). APOE traffics to astrocyte lipid droplets and modulates triglyceride saturation and droplet size. Journal of Cell Biology 223. 10.1083/jcb.202305003.

51. Cashikar, A.G., Toral-Rios, D., Timm, D., Romero, J., Strickland, M., Long, J.M., Han, X., Holtzman, D.M., and Paul, S.M. (2023). Regulation of astrocyte lipid metabolism and ApoE secretionby the microglial oxysterol, 25-hydroxycholesterol. J. Lipid Res. 64. 10.1016/J.JLR.2023.100350.

52. Farmer, B., Kluemper, J., and Johnson, L. (2019). Apolipoprotein E4 Alters Astrocyte Fatty Acid Metabolism and Lipid Droplet Formation. Cells 8, 182. 10.3390/cells8020182.

53. Garrahy, J.P. (2025). Astrocytic lipidopathy and bioenergetic failure in ApoE4-associated late-onset Alzheimer’s disease: A unifying hypothesis. Journal of Alzheimer’s Disease 106, 890–902. 10.1177/13872877251350338.

54. Cuní-López, C., Root, J.T., Hao, Y., Kowal, I., Blomberg, N., Ghirlando, R., Yang, L.G., Koppes-den Hertog, S.J., Cookson, M.R., van der Kant, R., et al. (2025). APOE genotypes differentially remodel the astrocytic lipid droplet-associated proteome to shape lipid droplet dynamics. bioRxiv. 10.1101/2025.08.19.669163.

55. Jacquemyn, J., Ralhan, I., and Ioannou, M.S. (2024). Driving factors of neuronal ferroptosis. Trends Cell Biol. 34, 535–546. 10.1016/j.tcb.2024.01.010.

56. Innerarity, T.L., Friedlander, E.J., Rall, S.C., Weisgraber, K.H., and Mahley, R.W. (1983). The receptor-binding domain of human apolipoprotein E. Binding of apolipoprotein E fragments. Journal of Biological Chemistry 258, 12341–12347. 10.1016/s0021-9258(17)44180-9.

57. Weisgraber, K.H., Innerarity, T.L., and Mahley, R.W. (1982). Abnormal lipoprotein receptor-binding activity of the human E apoprotein due to cysteine-arginine interchange at a single site. J. Biol. Chem. 257, 2518–2521. 10.1016/s0021-9258(18)34954-8.

58. Yamamoto, T., Choi, H.W., and Ryan, R.O. (2008). Apolipoprotein E isoform-specific binding to the low-density lipoprotein receptor. Anal. Biochem., 222–226.

59. Moon, H.J., Haroutunian, V., and Zhao, L. (2022). Human apolipoprotein E isoforms are differentially sialylated and the sialic acid moiety in ApoE2 attenuates ApoE2-Aβ interaction and Aβ fibrillation. Neurobiol. Dis. 164. 10.1016/j.nbd.2022.105631.

60. Poirier, S., Mayer, G., Benjannet, S., Bergeron, E., Marcinkiewicz, J., Nassoury, N., Mayer, H., Nimpf, J., Prat, A., and Seidah, N.G. (2008). The proprotein convertase PCSK9 induces the degradation of low density lipoprotein receptor (LDLR) and its closest family members VLDLR and ApoER2. Journal of Biological Chemistry 283, 2363–2372. 10.1074/jbc.M708098200.

61. Shan, L.X., Pang, L., Zhang, R., Murgolo, N.J., Lan, H., and Hedrick, J.A. (2008). PCSK9 binds to multiple receptors and can be functionally inhibited by an EGF-A peptide. Biochem. Biophys. Res. Commun. 375, 69–73. 10.1016/j.bbrc.2008.07.106.

62. Gu, H.M., Adijiang, A., Mah, M., and Zhang, D.W. (2013). Characterization of the role of EGF-A of low density lipoprotein receptor in PCSK9 binding. J. Lipid Res. 54, 3345–3357. 10.1194/jlr.M041129.

63. Xia, X.D., Peng, Z.S., Gu, H.M., Wang, M., Wang, G.Q., and Zhang, D.W. (2021). Regulation of PCSK9 Expression and Function: Mechanisms and Therapeutic Implications. Preprint at Frontiers Media S.A., 10.3389/fcvm.2021.764038 10.3389/fcvm.2021.764038.

64. Moulton, M.J., Barish, S., Ralhan, I., Chang, J., Goodman, L.D., Harland, J.G., Marcogliese, P.C., Johansson, J.O., Ioannou, M.S., and Bellen, H.J. (2021). Neuronal ROS-induced glial lipid droplet formation is altered by loss of Alzheimer’s disease-associated genes. Proceedings of the National Academy of Sciences 118. 10.1073/pnas.2112095118/-/DCSupplemental.

65. Canuel, M., Sun, X., Asselin, M.C., Paramithiotis, E., Prat, A., and Seidah, N.G. (2013). Proprotein Convertase Subtilisin/Kexin Type 9 (PCSK9) Can Mediate Degradation of the Low Density Lipoprotein Receptor-Related Protein 1 (LRP-1). PLoS One 8. 10.1371/journal.pone.0064145.

66. Stillman, J.M., Mendes Lopes, F., Lin, J.P., Hu, K., Reich, D.S., and Schafer, D.P. (2023). Lipofuscin-like autofluorescence within microglia and its impact on studying microglial engulfment. Nat. Commun. 14. 10.1038/s41467-023-42809-y.

67. Terman, A., and Brunk, U.T. (2004). Lipofuscin. Preprint at Elsevier Ltd, 10.1016/j.biocel.2003.08.009 10.1016/j.biocel.2003.08.009.

68. Zhou, X., Sun, L., Brady, O.A., Murphy, K.A., and Hu, F. (2017). Elevated TMEM106B levels exaggerate lipofuscin accumulation and lysosomal dysfunction in aged mice with progranulin deficiency. Acta Neuropathol. Commun. 5, 9. 10.1186/s40478-017-0412-1.

69. Moreno-García, A., Kun, A., Calero, O., Medina, M., and Calero, M. (2018). An overview of the role of lipofuscin in age-related neurodegeneration. Front. Neurosci. 12, 464. 10.3389/fnins.2018.00464.

70. Burns, J.C., Cotleur, B., Walther, D.M., Bajrami, B., Rubino, S.J., Wei, R., Franchimont, N., Cotman, S.L., Ransohoff, R.M., and Mingueneau, M. (2020). Differential accumulation of storage bodies with aging defines discrete subsets of microglia in the healthy brain. Elife 9, 1–71. 10.7554/eLife.57495.

71. Höhn, A., Jung, T., Grimm, S., and Grune, T. (2010). Lipofuscin-bound iron is a major intracellular source of oxidants: Role in senescent cells. Free Radic. Biol. Med. 48, 1100–1108. 10.1016/j.freeradbiomed.2010.01.030.

72. Martin, S., Harper, C.B., May, L.M., Coulson, E.J., Meunier, F.A., and Osborne, S.L. (2013). Inhibition of PIKfyve by YM-201636 Dysregulates Autophagy and Leads to Apoptosis-Independent Neuronal Cell Death. PLoS One 8. 10.1371/journal.pone.0060152.

73. Uwada, J., Nakazawa, H., Kiyoi, T., Yazawa, T., Muramatsu, I., and Masuoka, T. (2025). PIKFYVE inhibition induces endosome- and lysosome-derived vacuole enlargement via ammonium accumulation. J. Cell Sci. 138. 10.1242/jcs.262236.

74. Kutchukian, C., Casas, M., Dixon, R.E., and Dickson, E.J. (2025). Disruption of the PIKfyve complex unveils an adaptive mechanism to promote lysosomal repair and mitochondrial homeostasis. Nature Communications 16. 10.1038/s41467-025-65798-6.

75. Begum, G., Song, S., Wang, S., Zhao, H., Bhuiyan, M.I.H., Li, E., Nepomuceno, R., Ye, Q., Sun, M., Calderon, M.J., et al. (2018). Selective knockout of astrocytic Na+/H+ exchanger isoform 1 reduces astrogliosis, BBB damage, infarction, and improves neurological function after ischemic stroke. Glia 66, 126–144. 10.1002/glia.23232.

76. Siczkowski, M., and Ng, L.L. (1995). Culture density and the activity, abundance and phosphorylation of the Nat/H+ exchanger isoform 1ni human fibroblasts. Biochem. Biophys. Res. Commun.

77. Zhao, H., Carney, K.E., Falgoust, L., Pan, J.W., Sun, D., and Zhang, Z. (2016). Emerging roles of Na+/H+ exchangers in epilepsy and developmental brain disorders. Prog. Neurobiol. 138–140, 19–35. 10.1016/j.pneurobio.2016.02.002.

78. Marschallinger, J., Iram, T., Zardeneta, M., Lee, S.E., Lehallier, B., Haney, M.S., Pluvinage, J. V., Mathur, V., Hahn, O., Morgens, D.W., et al. (2020). Lipid-droplet-accumulating microglia represent a dysfunctional and proinflammatory state in the aging brain. Nat. Neurosci. 23, 194–208. 10.1038/s41593-019-0566-1.

79. Bae, J.Y., Jacquemyn, J., and Ioannou, M.S. (2024). Neuronal AMPK regulates lipid transport to microglia. Preprint at Elsevier Ltd, 10.1016/j.tcb.2024.08.001 10.1016/j.tcb.2024.08.001.

80. Li, Y., Munoz-Mayorga, D., Nie, Y., Kang, N., Tao, Y., Lagerwall, J., Pernaci, C., Curtin, G., Coufal, N.G., Mertens, J., et al. (2024). Microglial lipid droplet accumulation in tauopathy brain is regulated by neuronal AMPK. Cell Metab. 36, 1351–1370.e8. 10.1016/j.cmet.2024.03.014.

81. Tabor, G.T., Litvinchuk, A., Chen, Y., Allison, A., Sun, E.W., van Lengerich, B., Davis, S.S., West, E., Schlachetzki, J.C.M., Zeng, C., et al. (2026). Inducible deletion of DGAT1 and 2 from microglia exacerbates neurodegeneration and endolysosomal lipid accumulation in male PS19 mice. Cell Rep. 45, 116841. 10.1016/j.celrep.2025.116841.

82. Bhupana, J.N., Pabon, A., Leung, H.H., Rajmohamed, M.A., Kim, S.H., Tong, Y., Jang, M.H., and Wong, C.O. (2025). Endolysosomal processing of neuron-derived signaling lipids regulates autophagy and lipid droplet degradation in astrocytes. Nature Communications 16. 10.1038/s41467-025-60402-3.

83. Feringa, F.M., Koppes-den Hertog, S.J., Wang, L.Y., Derks, R.J.E., Kruijff, I., Erlebach, L., Heijneman, J., Miramontes, R., Pömpner, N., Blomberg, N., et al. (2025). The Neurolipid Atlas: a lipidomics resource for neurodegenerative diseases. Nat. Metab. 7, 2142–2164. 10.1038/s42255-025-01365-z.

84. Beretta, C., Dakhel, A., Eltom, K., Rosqvist, F., Uzoni, S., Mothes, T., Fletcher, J.S., Risérus, U., Sehlin, D., Rostami, J., et al. (2025). Astrocytic lipid droplets contain MHCII and may act as cogs in the antigen presentation machinery. Journal of Neuroinflammation 22. 10.1186/s12974-025-03452-0.

85. Gao, X., Gu, H., Li, G., Rye, K.A., and Zhang, D.W. (2012). Identification of an amino acid residue in ATP-binding cassette transport G1 critical for mediating cholesterol efflux. Biochim. Biophys. Acta Mol. Cell Biol. Lipids 1821, 552–559. 10.1016/j.bbalip.2011.07.012.

86. Hailey, D.W., Rambold, A.S., Satpute-Krishnan, P., Mitra, K., Sougrat, R., Kim, P.K., and Lippincott-Schwartz, J. (2010). Mitochondria Supply Membranes for Autophagosome Biogenesis during Starvation. Cell 141, 656–667. 10.1016/j.cell.2010.04.009.

87. Lord, S.J., Velle, K.B., Dyche Mullins, R., and Fritz-Laylin, L.K. (2020). SuperPlots: Communicating reproducibility and variability in cell biology. Preprint at Rockefeller University Press, 10.1083/JCB.202001064 10.1083/JCB.202001064.

